# Two-step mechanism of J-domain action in driving Hsp70 function

**DOI:** 10.1101/2020.01.13.901538

**Authors:** Bartłomiej Tomiczek, Wojciech Delewski, Lukasz Nierzwicki, Milena Stolarska, Igor Grochowina, Brenda Schilke, Rafal Dutkiewicz, Marta A. Uzarska, Szymon J. Ciesielski, Jacek Czub, Elizabeth A. Craig, Jaroslaw Marszalek

## Abstract

J-domain proteins (JDPs), obligatory Hsp70 cochaperones, play critical roles in protein homeostasis. They promote key allosteric transitions that stabilize Hsp70 interaction with substrate polypeptides upon hydrolysis of its bound ATP. Although a recent crystal structure revealed the physical mode of interaction between a J-domain and an Hsp70, the structural and dynamic consequences of J-domain action once bound and how Hsp70s discriminate among its multiple JDP partners remain enigmatic. We combined free energy simulations, biochemical assays and evolutionary analyses to address these issues. Our results indicate that the invariant aspartate of the J-domain perturbs a conserved intramolecular Hsp70 network of contacts that crosses domains. This perturbation leads to destabilization of the domain-domain interface - thereby promoting the allosteric transition that triggers ATP hydrolysis. While this mechanistic step is driven by conserved residues, evolutionarily variable residues are key to initial JDP/Hsp70 recognition - via electrostatic interactions between oppositely charged surfaces. We speculate that these variable residues allow an Hsp70 to discriminate amongst JDP partners, as many of them have coevolved. Together, our data points to a two-step mode of J-domain action, a recognition stage followed by a mechanistic stage.

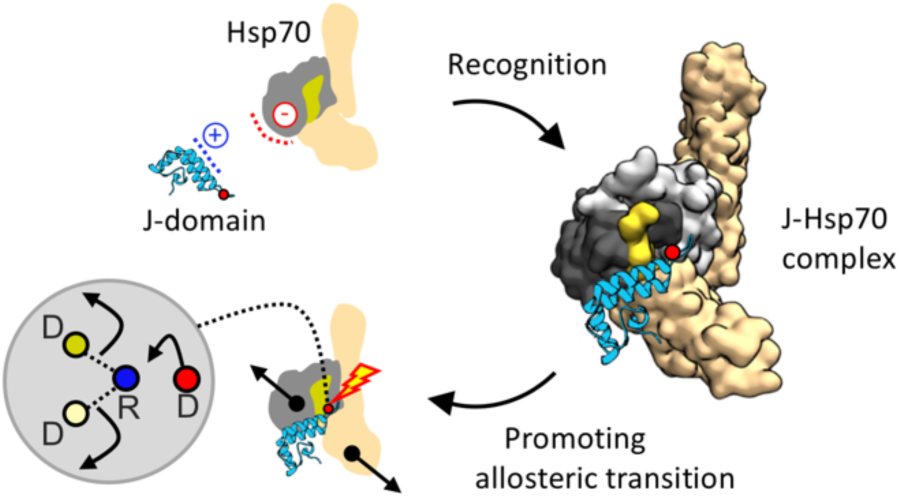

## Introduction

By transiently binding many different polypeptide substrates, Hsp70 chaperones assist in diverse cellular processes, from de novo protein folding to protein trafficking to disassembly of protein complexes. J-domain protein (JDP) cochaperones are crucial players in the cycle of interaction of Hsp70 with substrate. They transiently interact with ATP-bound Hsp70 via their defining J-domain. This interaction, in coordination with substrate binding, facilitates stimulation of Hsp70’s ATPase activity, which in turn drives the large scale conformational changes that stabilize substrate interaction^1-4^.

The structure of both J-domains and Hsp70s are conserved. J-domains are composed of four α-helices. Helices II and III form a finger like structure. The helix II/III connecting loop includes a conserved histidine, proline, aspartate tripeptide (HPD). These invariant residues are critical for J-domain stimulation of Hsp70’s ATPase activity. Hsp70s are more complex in structure, containing two large domains - a nucleotide binding domain (NBD) and a substrate binding domain (SBD) - connected by a “linker” segment. When ATP is bound, the domains are docked such that the substrate binding site in the β subdomain of the SBD (SBDβ) is easily accessible^5,6^. Upon ATP hydrolysis, the domains disengage, and the substrate is “trapped” as the α subdomain of the SBD (SBDα) closes over the SBDβ substrate binding site^7-9^.

The heart of the allosteric control mechanism behind these Hsp70 conformational transitions is an interaction network at the interdomain interface^10-12^. A critical segment of this interface is formed by interactions at the base of the NBD with the linker and SBDβ (called NBD/SBDβ,linker throughout)^5,6,13^. The linker plays an important role, as its interaction with the NBD is key to stimulation of ATPase activity^14-17^. A transient allosterically active intermediate poised for hydrolysis of ATP, often referred to as the “allosterically active state”, has been observed in the presence of a peptide substrate - NBD/SBD contacts are largely absent, but the linker remains bound to the NBD^10,18^.

It has been clear for years that J-domain interaction is essential for stimulation of Hsp70’s ATPase activity^19^. Particularly relevant to this report, early work on *Escherichia coli* JDP DnaJ and Hsp70 DnaK established an arginine at the Hsp70 NBD/SBDβ,linker interface, R167 of the NBD, as an important for J-domain function^14,20,21^. More recently a crystal structure of the J-domain of DnaJ in complex with ATP-bound DnaK was obtained ^22^. This normally transient interaction was trapped by covalently linking the J-domain and Hsp70 in cis, and by crosslinking Hsp70’s NBD and SBD to stabilize intramolecular interactions between domains. Consistent with the results of a wealth of mutational and biochemical analyses, as well as earlier molecular dynamics simulations^23^, the J-domain is bound at the NBD/SBDβ,linker interface. Residues of helices II and III and the connecting HPD loop form polar and hydrophobic interactions with the NBD and SBDβ.

Despite the accumulated information about J-domain/Hsp70 interactions, important questions remain. Although the end result of ATP hydrolysis is destabilization of Hsp70 domain-domain interactions, the initial structural consequences of J-domain binding are not known. Furthermore, there is no understanding of what determines the specificity of J-domain/Hsp70 interactions and how Hsp70s discriminate amongst their multiple JDP partners. To address these issues, we investigated two JDP/Hsp70 pairs: DnaJ/DnaK of *E. coli* and the mitochondrial Hsc20/Ssq1 of *Saccharomyces cerevisiae*^24,25^, which specializes in the biogenesis of proteins containing iron-sulfur clusters.

Results of our analyses point to two previously unappreciated facets of J-domain/Hsp70 interactions, leading us to propose a two-step model of J-domain action – recognition, followed by mechanistic action. Molecular simulations point to an active role of the invariant aspartate of the J-domain HPD in the mechanistic step – perturbing the intramolecular interaction network formed by the previously identified, key NBD arginine. This perturbation promotes allostery-related conformational changes of Hsp70. Evolutionary analyses indicate that J-domain/Hsp70 recognition, is dependent on variable, coevolving residues, even though the general mode of interaction of J-domains with Hsp70s is the same. This variability provides a plausible explanation for how Hsp70s are able to discriminate amongst their multiple JDP partners, thus enabling evolution of complex JDP/Hsp70 interaction networks.

## Results

### Structural model of the Hsc20-Ssq1 complex

To obtain a structural model of the JDP-Hsp70 complex formed by Hsc20 and Ssq1, we combined protein-protein docking with all-atom molecular dynamics (MD) simulations (Supplementary Fig. S1). This approach revealed a dominant bound state that accounted for 56% of the bound population (Fig. 1, Supplementary Fig. S2a). No other bound state accounted for more than 6% of the ensemble; these minor states rapidly interconverted and had a clear tendency to converge to the most populated bound state (Supplementary Fig. S2b). The binding mode in this dominant state closely resembles the recently published X-ray structure of DnaJ’s J-domain in complex with DnaK (DnaJ^JD^-DnaK)^22^. Helices II and III are oriented very similarly in the two complexes - with a backbone root mean square deviation (RMSD) of 4.17 Å (Supplementary Fig. S3). More specifically, the J-domains bind at the NBD/SBDβ,linker interface such that helix II predominantly interacts with the β-sheet region of NBD subdomain IIa and helix III with SBDβ (Fig. 1). The HPD residues are positioned nearly identically in the two complexes, towards R167/R207 of NBD subdomain Ia in DnaK/Ssq1 (referred to as R^NBD^ throughout), the arginine that previous analyses of DnaK established as important for allosteric transitions^14,20,21^.

**Figure 1.**
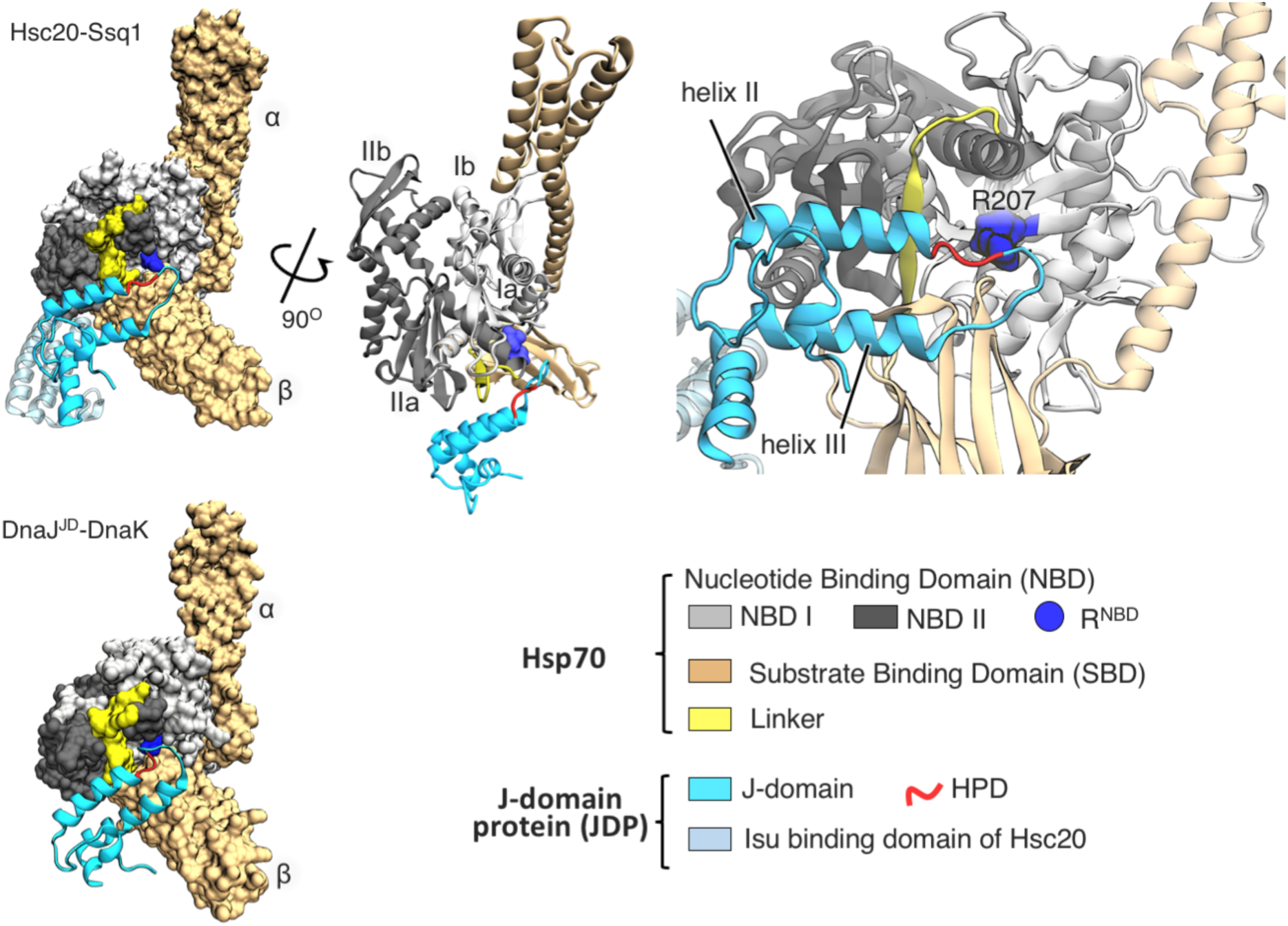
Binding mode of J-domain-Hsp70 complexes. (*top left*) The dominant state of the Hsc20-Ssq1 complex obtained by protein-protein docking followed by MD refinement. Structure of *S. cerevisiae* Hsc20 (PDB ID 3UO3) was used along with a homology model of ATP-bound Ssq1, based on the X-ray structure of ATP-bound DnaK (PDB ID 4B9Q). The Ssq1 model had the expected architecture of an ATP-bound Hsp70: Nucleotide binding domain (NBD, grey) interacting with α and β subdomains of the substrate binding domain (SBD, brown) and the linker (Lk, yellow), in a groove between NBD subdomains Ia (light grey) and IIa (dark grey). To model the Hsc20-Ssq1 complex, conformational fluctuations of Hsc20 and ATP-bound Ssq1 were first characterized using MD simulations and clustering (Supplementary Fig. S1). The obtained distinct conformers were then combinatorially docked. Based on the energetic and geometric criteria described in Methods, a representative set of 33 of the resulting complexes was selected for further refinement with MD simulations, and then clustered according to the positioning of the J-domain with respect to Ssq1 (Supplementary Fig. S2a). The model shown represents the dominant state with a population of 56%: J-domain of Hsc20 (cyan) interacts with Ssq1 at the NBD/SBDβ, linker interface. HPD sequence (red) of the J-domain is positioned near the conserved R207 (blue) of the NBD. (*top middle*) View of the Hsc20-Ssq1 complex to illustrate the orientation of the J-domain relative to the NBD subdomains (IA, IB, IIA, IIB s) which are marked, only the J-domain of Hsc20 is shown for clarity. (*top right*) Close-up showing J-domain interaction with an orientation similar to that at left. (*bottom left*) X-ray structure of the J-domain of DnaJ (DnaJ^JD^) in complex with DnaK (DnaJ^JD^-DnaK) (PDB ID 5NRO).

### D^HPD^ interferes in Hsp70 interdomain interactions by perturbing critical R^NBD^ contacts

To address the question of the mechanism of J-domain action, we employed MD simulations that, unlike most other methods, allow studying of transient residue-residue interactions^26^. We first analyzed Hsp70 alone, starting with the Ssq1 structural model - focusing on intramolecular interactions involving the R^NBD^ because of its demonstrated importance in NBD/SBD communication^14,20,21^ and its proximity to the HPD in the J-domain-Hsp70 complexes. Our initial analysis of MD trajectories revealed R^NBD^ to be capable of forming two ion pairs, one with D517 of the SBDβ (referred to as D^SBD^ throughout) and one with D429 of the interdomain linker (referred to as D^Lk^ throughout). To analyze these two contacts of R^NBD^ more closely, we constructed the free energy map governing their formation (Fig. 2a), by applying metadynamics, an enhanced sampling method suitable for simulating complex conformational changes^27,28^. As relevant coordinates, we used the distances between R^NBD^ and its interaction partners D^SBD^ and D^Lk^. We found the probability of the two contacts occurring at the same time to be 82%. In the DnaK crystal structure^5^ the homologous D^SBD^-R^NBD^ pair is present, but the homologous D^Lk^-R^NBD^ pair is not. To address this apparent discrepancy, we performed metadynamics simulations of the DnaK crystal structure. Analysis of the free energy map revealed that DnaK R^NBD^ interacts simultaneously with D^SBD^ and D^Lk^ 83% of the time. Such high frequencies suggest that, though not captured in the static DnaK crystal structure, this arginine double interaction occurs frequently in both Ssq1 and DnaK.

**Figure 2.**
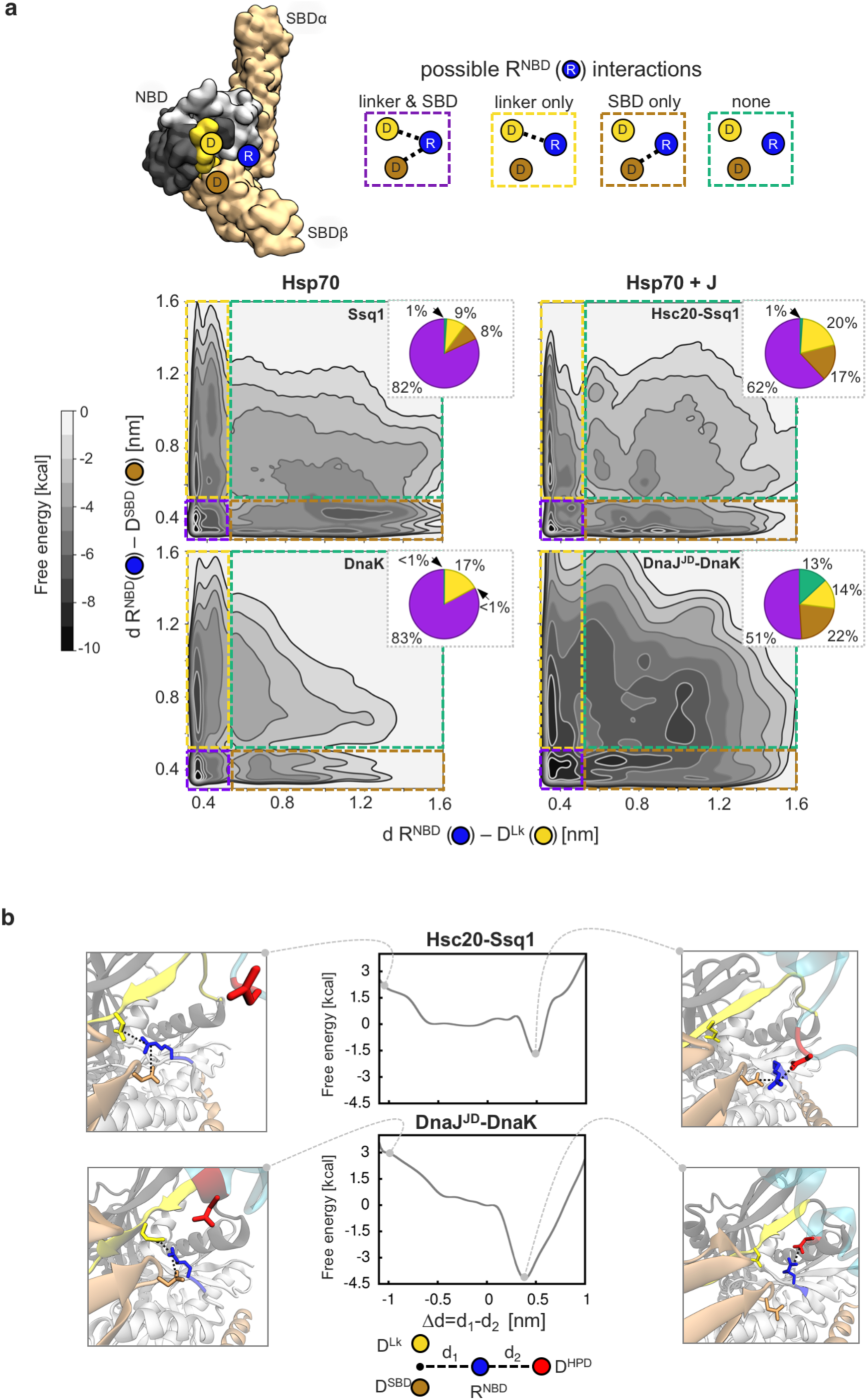
D^HPD^ of J-domain displaces interdomain contacts formed by R^NBD^ with the D^Lk^ and D^SBD^. (a) Free energy landscape governing the formation of Ssq1/DnaK R^NBD^ interdomain contacts with D^Lk^ and D^SBD^ (*top*) Schematic depicting of the three Hsp70 residues of the R^NBD^ interaction network; R^NBD^ of subdomain Ia (blue dot) can interact with D^LK^ of the linker (yellow dot) and D^SBD^ of the SBDβ (brown dot). The four possible interaction states of R^NBD^ are illustrated at right with the color of the rectangles serving as a key for the free energy landscapes at bottom. NBD subdomains Ia (light grey) and IIa (dark grey), SBD subdomains α and β (brown), linker (yellow). R^NBD^ = R207/R167 in Ssq1/DnaK; D^SBD^ = D429/D393 in Ssq1/DnaK; D^Lk^ = D517/D481 in Ssq1/DnaK. (*bottom*) Free energy landscapes governing the formation of these interaction states for Ssq1 and DnaK alone (left panels) and for Hsc20-Ssq1 and DnaJ^JD^-DnaK complexes (right panels) with distances between R^NBD^ -D^Lk^ and between R^NBD^-D^SBD^ used as relevant coordinates. Integrated populations of these interaction states are shown by pie charts. (b) Free energy profile of the transition of R^NBD^ from intramolecular interactions with D^Lk^ and D^SBD^ residues to intermolecular contact with D^HPD^ of the J-domain HPD loop when kept near NBD subdomain Ia. Distance difference Δd = d_1_ - d_2_ is used as a relevant coordinate. d_1_ and d_2_ are the distances between R^NBD^ and D^Lk^/D^SBD^ of Hsp70 (d1) or D^HPD^ of the J-domain (d_2_). Conformations in which R^NBD^ interacts with D^HPD^ at the expense of interaction with D^Lk^ and/or D^SBD^ (representative structures on the right, corresponding to the free energy minimum at 0.4-0.5 nm) are energetically more favorable than conformations with R^NBD^ interacting simultaneously with D^Lk^ and D^SBD^ (representative structures on the left).

To assess the effect of J-domain binding to Hsp70 on this conserved arginine-centered interaction network, we carried out the same set of calculations for Ssq1 and DnaK in complex with Hsc20 and DnaJ^JD^, respectively (Fig. 2a). For both Hsc20-Ssq1 and DnaJ^JD^-DnaK, the fraction of the population of Hsp70 molecules in which R^NBD^ contacts both D^SBD^ and D^LK^ was reduced compared to that of Hsp70 alone: Ssq1, from 82 to 62%; DnaK, from 83 to 51%. We hypothesized that D^HPD^ competes with D^SBD^ and D^Lk^ for interactions with R^NBD^ and therefore is responsible for the observed partial loss of its intramolecular contacts. To determine the likelihood of such an exchange of interaction partners, we performed metadynamics simulations to compute the free energy profiles for transition of R^NBD^ from intramolecular interaction with D^SBD^ and D^Lk^ to interaction with D^HPD^ (Fig. 2b). In these simulations, the HPD loop was kept near NBD subdomain Ia in the orientation predisposing it for the interaction with R^NBD^ (see Fig. 1). The well-defined free energy minima in the resulting profiles indicate that R^NBD^ shows preference (1.5 and 4 kcal/mol for Ssq1 and DnaK, respectively) for interaction with D^HPD^, rather than simultaneous interaction with the aspartates on the SBD and the linker, D^SBD^ and D^Lk^. An example of spontaneous transition of R^NBD^ to the intermolecular contact with D^HPD^, captured in our unbiased equilibrium MD simulation of the Hsc20-Ssq1 complex, is shown in Movie S1. Taken together, our data suggest that D^HPD^ perturbs the R^NBD^ -centered interaction network at the NBD/SBDβ,linker interface.

### J-domain binding initiates SBDβ disengagement from the NBD

We next asked how perturbation of intramolecular interactions of R^NBD^ with D^SBD^ and D^Lk^ affects the NBD/SBDβ,linker interface. First, we considered the effect of substitution of R^NBD^, such that interactions with aspartates are prohibited. An R^NBD^->D DnaK substitution variant has been shown to be defective in interdomain communication^14^. Its intrinsic ATPase activity is somewhat higher than that of wild-type (WT) DnaK and not effectively stimulated in the simultaneous presence of its JDP partner DnaJ and protein substrate. We therefore asked whether substitution of R^NBD^->D in Ssq1 had a similar effect. We found the basal ATPase activity of Ssq1 R^NBD^->D to be 3-fold higher than that of WT Ssq1, and not efficiently stimulated in the presence of Hsc20 and protein substrate, reaching only 30% of the activity observed for WT Ssq1 (Supplementary Fig. S4). Thus, substitution of the conserved R^NBD^ has significant effects on interdomain communication in both DnaK and Ssq1.

Because of these effects on interdomain communication, we used MD simulations to probe how weakening of the cross-domain interactions of R^NBD^ (i.e. with D^SBD^ and D^Lk^) affects the compactness of the NBD/SBDβ,linker interface. Umbrella sampling was used because it allows testing of conformations not accessible by conventional MD simulations. For these simulations a center of mass distance between SBDβ and its NBD interacting region (SBDβ-NBD distance) was used as a relevant coordinate. To accelerate the free energy convergence and to allow exhaustive sampling of interdomain contacts these simulations were performed using Ssq1 and DnaK with a truncated SBDα subdomains.

Free energy profiles were computed (Supplementary Fig. S5), and then converted into probability distributions (Fig. 3a). These distributions show that Ssq1 and DnaK both adopt conformations with longer SBDβ-NBD distances in the presence, than in the absence of a J-domain, indicative of a transition to a less compact interdomain interface. Furthermore, the probability distributions of the R^NBD^->D variants (Fig. 3a) resemble those of Hsp70s in the presence of the J-domain in that that the mean distance values are very similar for both distributions, supporting our hypothesis that disruption of the arginine centered interaction network affects compactness of the interdomain interface.

**Figure 3.**
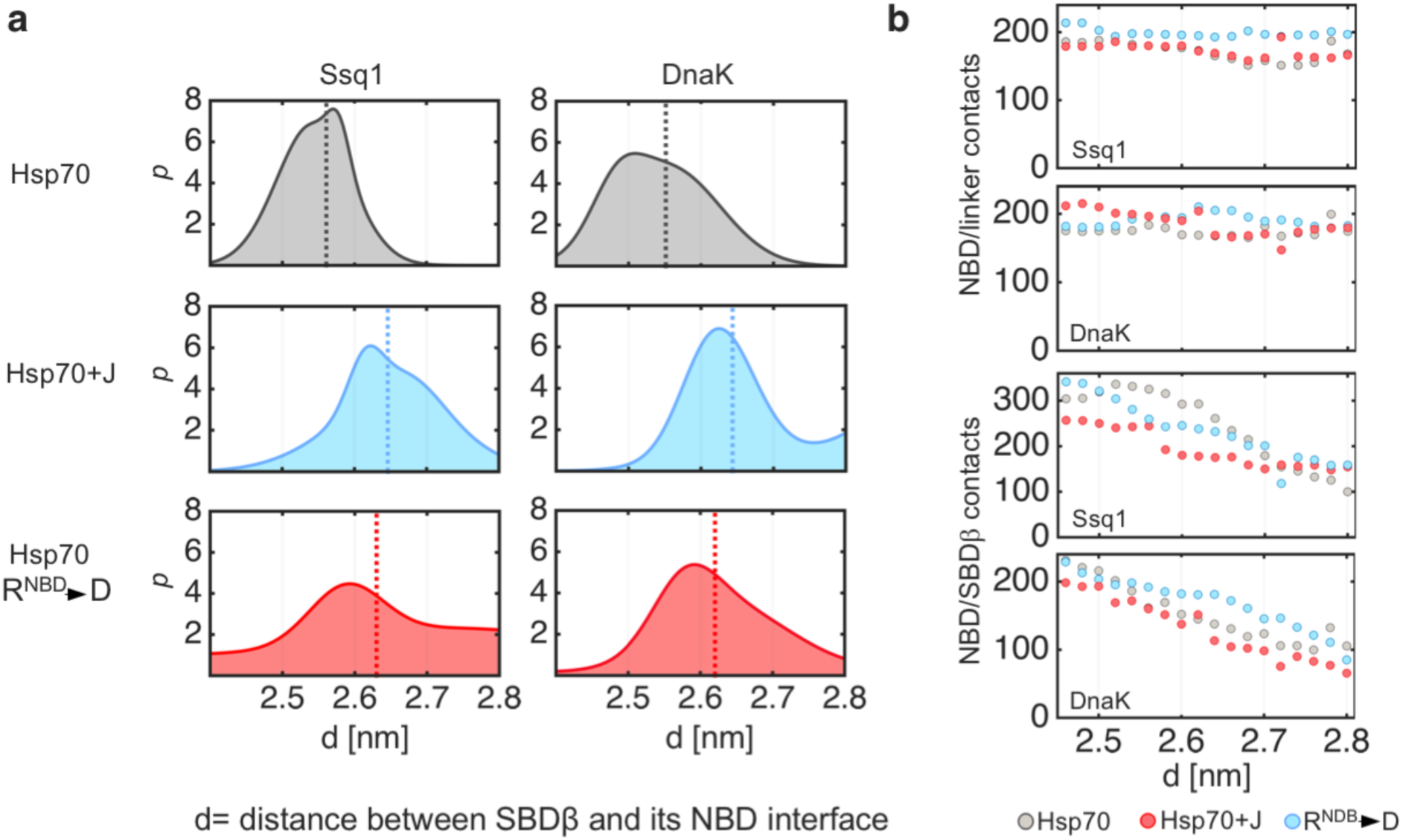
J-domain binding displaces SBDβ from association with NBD. (a) Probability distributions of the center of mass distance d between SBDβ and its NBD interface for Ssq1 and DnaK; Ssq1 and DnaK alone (top panels, in grey), Hsc20-Ssq1 and DnaJ^JD^-DnaK complexes (center panels, in cyan), R^NBD^ substitution variants Ssq1(R207D) and DnaK(R167D) (bottom panels, in red). Dashed lines represent an average d value for each distribution. (b) The number of contacts between NBD and linker and between NBD and SBDβ, with increasing distance d; Ssq1 and DnaK (grey dots); Hsc20-Ssq1 and DnaJ^JD^-DnaK (cyan dots); Ssq1(R207D) and DnaK(R167D) (red dots). Contacts were defined as the number of non-hydrogen atoms within a 0.5 nm cutoff distance across the binding interface.

To assess what structural changes took place within the NBD/SBDβ,linker interface we used umbrella sampling trajectories to determine the number of interdomain contacts (Fig. 3b). We analyzed interactions between the NBD and SBDβ and between the NBD and the linker separately, because of their opposing effects on ATPase activity (i.e. ATPase activity requires linker association and SBD dissociation). In all our analyses, for both Ssq1 and DnaK, the number of the NBD/linker contacts remains virtually the same - regardless of the presence of the J-domain or the R^NBD^->D substitution. In contrast, the number of NBD/SBDβ contacts were fewer (by up to 50 %) in the presence of the J-domains or the R^NBD^->D substitution, indicating that perturbation of the arginine-centered interaction network promotes partial disengagement of SBDβ. Taken together, these results indicate that J-domain binding perturbs the R^NBD^ intramolecular interactions which in turn leads to partial disengagement of SBDβ, while the interdomain linker remains tightly associated with the NBD. Such an arrangement favors ATP hydrolysis, which triggers further conformational changes^10,11,14,15^.

### Conservation of residues involved in the mechanistic step of J-domain action

The R^NBD^-centered interaction network is the same in DnaK and Ssq1, in that R^NBD^ interacts with homologous aspartates in the linker and SBDβ. To investigate the prevalence of these residues, we analyzed Hsp70 sequences from a large number of bacterial and eukaryotic species. R^NBD^ and its interaction partner D^Lk^ is invariant in our data set (Table 1). The conservation of its SBDβ interaction partner D^SBD^ is somewhat more complex. Aspartate is only present at this position in DnaKs of bacteria closely related to *E. coli* and in mitochondrial Hsp70s of fungi; asparagine is present at this position in most Hsp70s (Table 1, Supplementary Fig. S6). That asparagine can also form a hydrogen bond with arginine^29^, thereby participating in the interaction network, supports the idea that the arginine-centered network is conserved across Hsp70s. Thus, it is likely the mechanism of action of J-domains, that is perturbation of this network, is conserved as well.

**Table 1.** Evolutionary conservation of R^NBD^-centered interaction network.

|  | number of sequences | residue conservation |  |  |
| --- | --- | --- | --- | --- |
|  |  | R<sup>NBD</sup> | D<sup>SBD</sup> | D<sup>Lk</sup> |
| mitochondrial Hsp70 | 404 | 100% | 100% | 33% D (65% N) |
| DnaK | 1560 | 100% | 100% | 17% D (83% N) |
| cytosolic Hsp70 | 586 | 100% | 100% | 2% D (98% N) |
| ER Hsp70 | 334 | 100% | 100% | 2% D (98% N) |

### Importance of electrostatic interactions at the Hsc20/Ssq1 binding interface

Having found similarities of the R^NBD^ network and the overall mode of interaction of DnaJ^JD^-DnaK and Hsc20-Ssq1, we decided to look more closely at the J-domain/Hsp70 binding interfaces. DnaJ^JD^-DnaK has been studied extensively^22^. However, no information is available for Hsc20-Ssq1. We therefore expanded our computational analysis of the Hsc20-Ssq1 complex. We used MD simulation trajectories to search for all specific (polar and hydrophobic) residue-residue contacts, and computed the relative energetic contributions of individual residues to complex stability (Supplementary Fig. S7-8). The binding interface is formed by 17 residues of the J-domain contacting 18 residues of Ssq1 at a cutoff distance of 0.5 nm. The interface, which occupies 6.7 ± 0.2 nm^2^, is mostly hydrophilic, with scattered hydrophobic interactions contributing 12.5 % of the surface (0.8 ± 0.1 nm^2^). The predominant contributors are eight charged residues (Fig. 4a). Three positively charged residues of Hsc20 helix II (R37^JD^, K38^JD^, R41^JD^) that form a network of electrostatic interactions with four negatively charged residues of the Ssq1 NBD (D246, E248, D249, E253) and K70^JD^ of helix III that promotes complex formation, though its binding partner(s) was not identified in our analyses.

**Figure 4.**
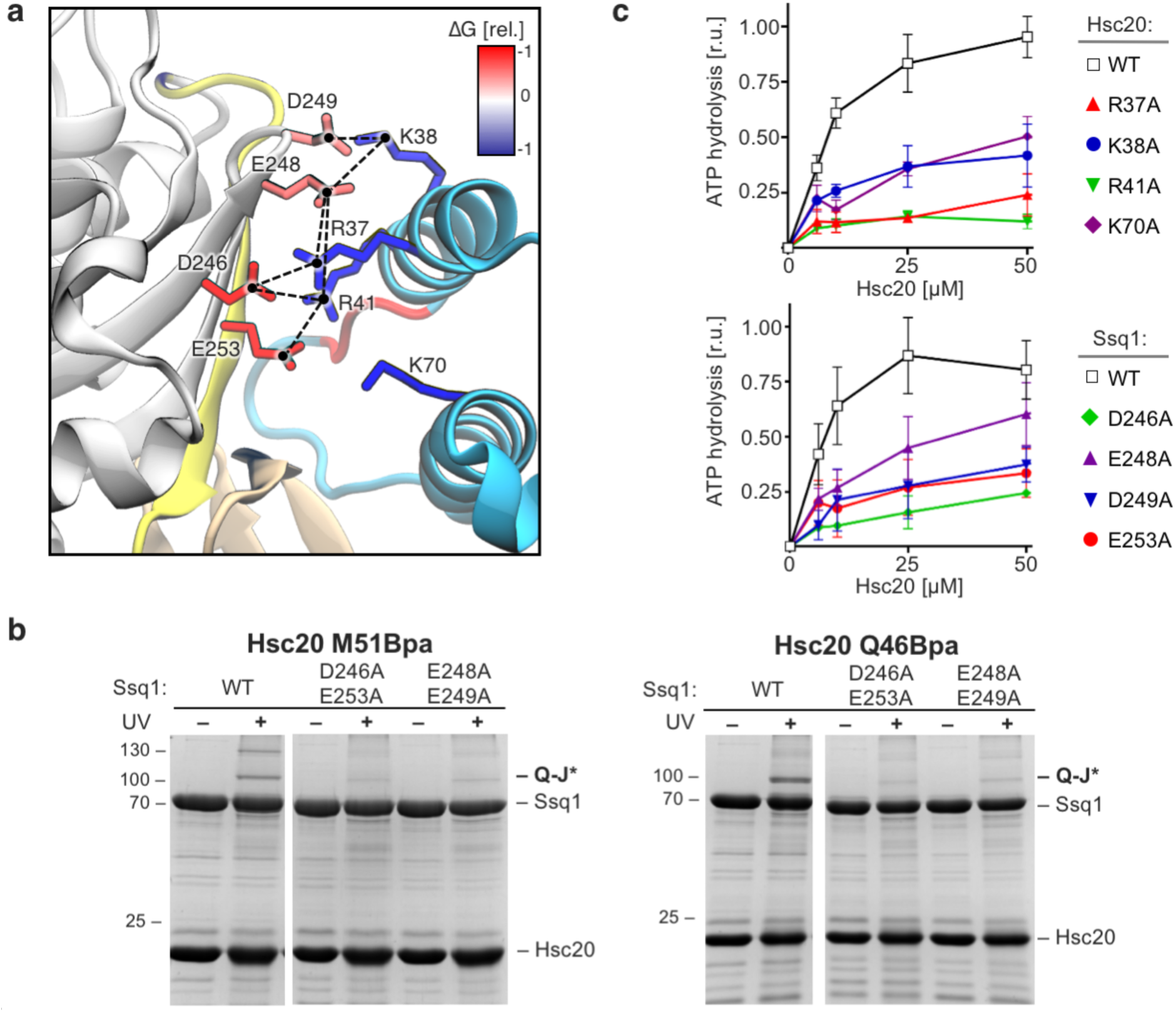
Electrostatic interface of Hsc20 J-domain and Ssq1. (a) Network of key polar interactions at the binding interface between the J-domain of Hsc20 (cyan) and Ssq1 subdomain IIa (grey), linker (yellow) and SBDβ (brown). The most stable ion pairs and hydrogen bonds identified in MD simulations are indicated with dashed lines. Relative contributions of individual residues to the Hsc20-Ssq1 complex stability are indicated by color scale; blue for J-domain residues and red for Ssq1 residues. Note: K70^JD^ does not form obvious contacts but contributes strongly to the complex stability. (b) Site specific crosslinking of Hsc20 to Ssq1. Purified Hsc20 Q46Bpa (left) and M51Bpa (right) variants were incubated with purified Ssq1 wild-type (WT) or alanine substitution variants (D246A, E253A or E248A, D249A) in the presence of ATP. After UV irradiation (+), or as a control no irradiation (-), reaction mixtures were separated by SDS-PAGE. Migrations of size standard, in kDa, are indicated; Q-J* indicates position of the Ssq1-Hsc20 crosslink, which was identified using mass-spectroscopy (Supplementary Fig. S9). (c) Stimulation of Ssq1 ATPase activity by Hsc20 was measured in the presence of 0.5 μM of Ssq1, 500 μM of PVK-peptide substrate, 0.5 μM of nucleotide exchange factor Mge1 and indicated concentrations of Hsc20. (top) Hsc20 WT and alanine substitution variants. (bottom) Ssq1 WT and alanine substitution variants. The ATPase activity corresponding to maximal stimulation of the WT Hsc20 was set to 1; error bars indicate standard deviation for three independent measurements.

We carried out two types of biochemical experiments to verify the computationally derived model of the Hsc20-Ssq1 complex. First, we asked if alanine substitution of these Ssq1 residues disturbs the Hsc20/Ssq1 interaction. To enable detection of the inherently transient J-domain/Hsp70 interaction, we used Hsc20 having a site-specific p-benzoyl-l-phenylalanine (Bpa) crosslinker^30^ replacing two residues, one upstream (Q46) and one downstream (M51) of the HPD (Fig. 4b, Supplementary Fig. S9). The efficiency of Hsc20 crosslinking to the Ssq1 variants was strongly reduced compared to that of WT Ssq1. Second, we measured Hsc20’s ability to stimulate the ATPase activity of Ssq1. We tested four Hsc20 and four Ssq1 variants, each having a different single alanine substitution. All variants had reduced activity (Fig. 4c). We conclude that the key residues of Hsc20 and Ssq1 identified by the MD simulations are indeed important for the Hsc20/Ssq1 interaction.

### Hsc20 binding to Ssq1 is driven by long-range electrostatic interactions

The importance of charged residues in Hsc20-Ssq1 complex stability led us to consider whether the initial encounter of the two proteins is electrostatically driven - an idea furthered by the presence of a patch of uniformly positive electrostatic potential around helix II of the J-domain and a complementary patch of negative potential around subdomain IIa of the Ssq1 NBD, the site of J-domain interaction (Supplementary Fig. S10a). MD simulations in which Hsc20 and Ssq1 were initially separated with their charged patches facing each other, revealed that Hsc20 spontaneously interacted with Ssq1 within tens of ns and remained stably bound for the rest of the stimulation (Movie S2). Notably, the spontaneously formed complexes are very similar to the dominant bound mode identified in our systematic search (Supplementary fig. S10c).

To examine the driving forces behind spontaneous Hsc20/Ssq1 interaction, we computed the underlying free energy profile using umbrella sampling. The resulting profile (Supplementary Fig. S10d, shows a pronounced bound-state energy minimum (at distances < 2 nm), indicative of site-specific binding. The non-zero slope of the WT free energy profile up to ∼3.5 nm demonstrates that Hsc20 is attracted to Ssq1 by long-range electrostatic driving forces. Consistent with the above analysis of the binding interface and biochemical data (Fig. 4), the free energy minimum, and hence binding affinity, almost completely disappears in the case of the Hsc20 variant having helix II alanine substitutions R37A,R41A (Supplementary Fig. S10b,d).

### Residues forming the JDP/Hsp70 interface are variable and coevolving

The two divergent J-domain/Hsp70 partnerships discussed here show remarkable structural similarity in their overall mode of interaction - helix II with the β-sheet of NBD subdomain IIa and helix III with SBDβ (Fig. 1). However, when we aligned these regions it became apparent that the residues forming the two J-domain/Hsp70 binding interfaces are not conserved (Fig. 5a). This is particularly evident for the J-domains - only 12 of the 17 positions forming the Hsc20/Ssq1 interface are shared with the DnaJ^JD^/DnaK interface, and of these 12, only 3 are occupied by the same amino acids in the two interfaces. A broader comparison of Hsc20, Ssq1, DnaJ, and DnaK with their orthologs from a wide range of species belonging to fungi, animals and plants (Supplementary Fig. S11-12) revealed that the amino acids occupying the J-domain/Hsp70 interfacial positions are indeed variable. The J-domain side of these interfaces is particularly so. Variability is evident for J-domains of Hsc20 and DnaJ among relatively closely related species, Ascomycota and Bacteria, respectively, suggesting that even closely related JDP/Hsp70 pairs engage different sets of residues to form the binding interface (Fig. 5b). One trend is evident however. Despite the overall variability, the J-domain side of the interface is enriched in positively charged residues (Fig. 5c) - consistent with our results that complementary electrostatic interactions are crucial for stability of the Hsc20-Ssq1 complex. The Hsp70 side of the binding interface is less variable than the J-domain side; 8 out of the 16 positions shared between the Ssq1/DnaK interfaces are occupied by identical amino acids.

**Figure 5.**
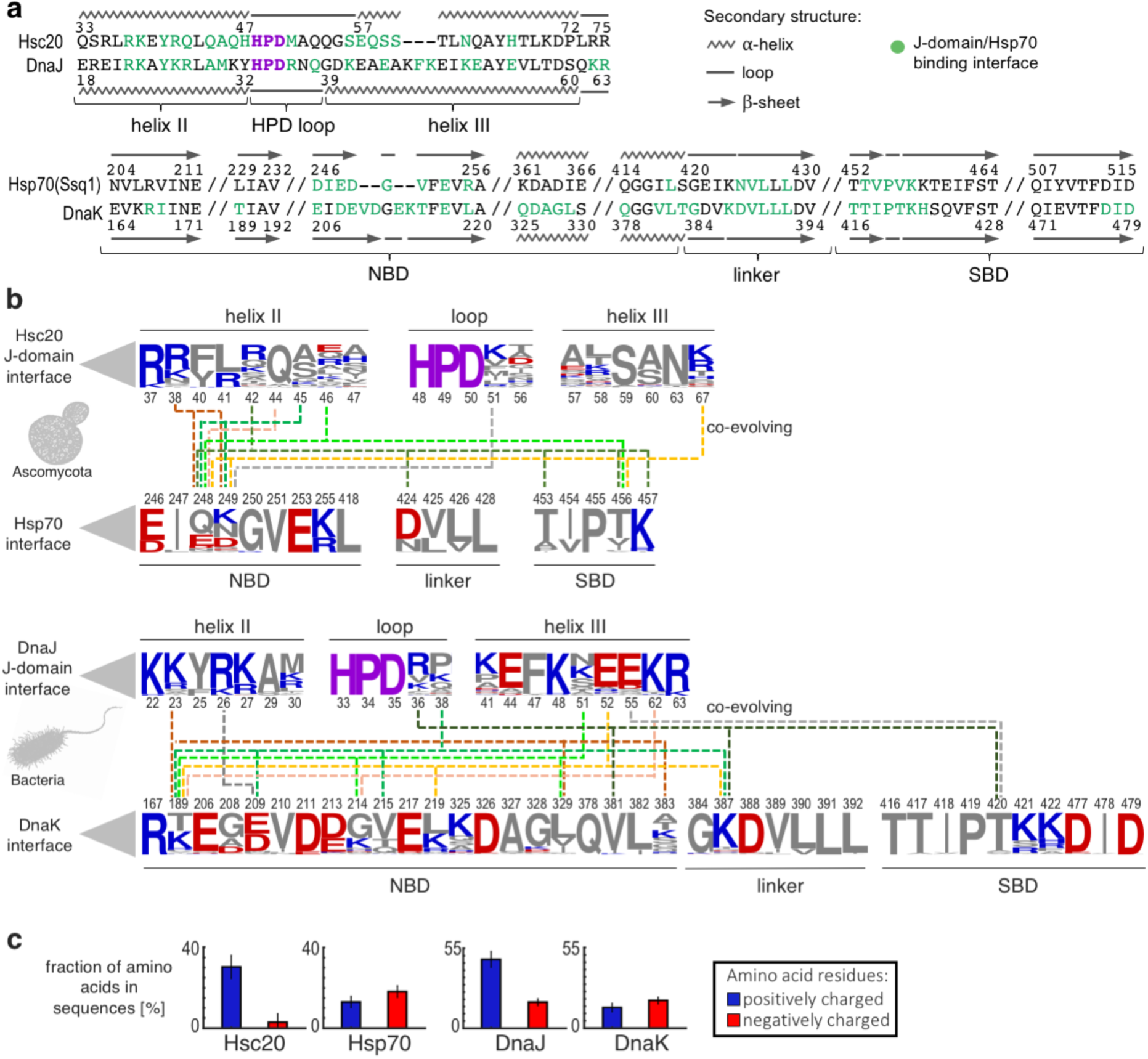
Sequence variability of J-domain/Hsp70 binding interface. (a) Pairwise sequence alignments of the J-domain/Hsp70 interfacial regions of the J-domains Hsc20 and DnaJ’s (top) and the Hsp70s Ssq1 and DnaK (bottom). Residues of each protein that are in contact across the Hsc20/Ssq1 and DnaJ^JD^/DnaK binding interfaces are in green; HPD motifs in purple. (b) Sequence variability of positions that form the J-domain/Hsp70 binding interfaces (those positions shown in green in panel a) for orthologs of Hsc20/Ssq1 from 67 Ascomycota species (top) and DnaJ/DnaK from 117 bacteria species (bottom). Coevolving positions with p>0.01 statistical support (Supplementary Table S3-4, Fig. S16-17) are connected by dashed lines. Sequence logos represent the amino acid frequency of each interfacial position in the orthologs examined, with positively charged (blue), negatively charged (red) and uncharged (grey) residues. Position numbering is that for *S. cerevisiae* (top) and *E. coli* (bottom) sequences. (c) Fraction of interfacial Hsc20/Ssq1 and DnaJ^JD^/DnaK residues that are positively and negatively charged. Whiskers indicate standard deviation.

The data discussed in the previous paragraph presents an apparent conundrum – variability of J-domain/Hsp70 interfacial residues, yet maintenance of the functional interaction of J-domains with Hsp70 partners. Coevolution of residues across the binding interface, such that substitutions on one side are compensated by complementing substitutions on the other^31^, could be an explanation. Therefore, we tested whether J-domain/Hsp70 interfacial residues are coevolving. We analyzed the Ascomycota and bacterial data sets discussed above (Fig. 5b), applying the probabilistic Coev model^32^ of sequence evolution that allowed us to discriminate between coevolving and independently evolving positions, while incorporating the underlying phylogenetic relationships (Supplementary Fig. S13-14). We found that some of the variable positions are indeed coevolving - 16 for the Hsc20/Hsp70 interface; 20 for DnaJ/DnaK (Fig. 5b). As a control, we tested the same datasets using a model of ‘independent’ sequence evolution. The Coev model fits our data significantly better than the independent model (Supplementary Table S2-3 and Fig. S15-16), supporting our hypothesis that coevolution of interacting partners could explain how J-domain/Hsp70 functional interactions are maintained despite sequence diversity.

## Discussion

Based on the results presented we propose a mechanism of J-domain function that reconciles two apparently contradictory features: an invariant dependence on the HPD signature motif, yet an ability to recognize a specific Hsp70 partner. Our data supports a model in which the D^HPD^ of the J-domain plays an active role in perturbing the intramolecular interactions of the Hsp70 R^NBD^ with conserved residues of the linker and SBDβ, destabilizing NBD/SBD interactions and thus promoting allosteric transition leading to ATP hydrolysis. On the other hand, evolutionarily variable J-domain residues are responsible for the specificity of exclusive JDP/Hsp70 interactions, as well as fine-tuning prioritization of JDP/Hsp70 interactions in the cases where an Hsp70 has multiple JDP partners that guide Hsp70 to different cellular functions.

### Mechanistic step

Based on detection of an energetically favorable interaction between invariant D^HPD^ and invariant R^NBD^, we propose that the D^HPD^ -R^NBD^ interaction displaces those of R^NBD^ with residues of the interdomain linker (D^Lk^) and SBDβ subdomain (D^SBD^) of Hsp70. This displacement promotes destabilization of the interdomain interface, such that the SBDβ partially disengages from the NBD. It is important to note that the interdomain linker remains stably bound to the NBD upon J-domain induced disengagement of the SBDβ. This structural arrangement has been termed the “allosterically active state” ^10^ because linker engagement with, but SBDβ disengagement from, the NBD is necessary for ATPase activation that triggers further Hsp70 conformational changes required for stabilization of substrate binding^11,14-17,33^. The data presented here also allows a more unified picture of how Hsp70 reaches the allosterically active state to effectively trap substrate. It was previously shown that the allosterically active state is populated in the presence of ATP and substrate^10,11,34^. Our data indicates that it is also populated in the presence of ATP and J-domain. Thus, the substrate and J-domain dependent allosteric pathways converge. In the case of substrate binding an allosteric signal is propagated through a cascade of interactions from the SBDβ substrate binding site to the NBD/SBDβ,linker interface^10-12,34^; in the case of J-domain binding the signal is propagated more directly by perturbation of the R^NBD^ centered interaction network at the interface. We suggest that this convergence allows synchronization - only upon simultaneous interaction of substrate and J-domain is ATPase activity effectively stimulated, driving transition to the undocked state and substrate capture.

This coordinated mechanism also impacts the larger picture of Hsp70 allosteric dynamics. All Hsp70s studied share major characteristic transient conformational shifts – for example, undergoing repeated transitions involving undocking from the NBD of both SBDβ and SBDα. However, recent single molecule experiments indicate that the frequency of such transitions differs among members of the Hsp70 family^7-9,18,35-37^. Interestingly, eukaryotic Hsp70s such as BiP of the endoplasmic reticulum^8,36^ or Hsp70 and Hsc70^37^ of the cytosol, display conformational dynamics markedly different from DnaK and mitochondrial Hsp70. They have D^SBD^ replaced by asparagine, pointing to the importance of the R^NBD^ centered interactions for the allostery of Hsp70s.

Though no model for the mechanistic role of the D^HPD^-R^NBD^ interaction per se has been previously put forth, the functional connection between these residues is well supported by genetic and biochemical data. Alteration of either R^NBD^ or residues interacting with it (D^SBD^ and D^Lk^)^10,11,14,17,34,38^disrupt allosteric communication between domains, resulting in a somewhat increased basal activity and ineffective stimulation by its JDP partner, DnaJ in the case of DnaK^14^ and Hsc20 in the case of Ssc1 (Supplementary Fig. S4). Substitution of R^NBD^(R167H) of DnaK suppresses nonfunctional D^HPD^(D35N) variant of DnaJ^20^. However, we do not mean to say that this role of the D^HPD^ is the only mechanistic effect of J-domain binding to Hsp70. For example, the invariant histidine and/or proline of the HPD may play specific roles, and, as previously suggested^39^, conformational strain induced in the J-domain upon binding to Hsp70 might be important as well. However, there is no question that R^NBD^ is key and could be considered as an allosteric perturbation site for the J-domain action.

### Recognition step

At first glance, the sequence diversity of the interface of J-domain-Hsp70 complexes, but maintenance of overall positioning, and thus, function, seems surprising. It raises the question of how this diversity evolved without loss of function. Our analyses suggest that functional J-domain/Hsp70 interactions have been maintained over time via coevolution of residues that form the binding interface. The nature of the Hsc20/Ssq1 interface is instructive in this regard. Each key residue on one side of the large interface contacts more than one residue on the opposite side. Such a multifaceted interaction network is highly flexible. Over time residues can be added to or deleted from the interface without ever losing the ability to maintain functional interaction between the coevolving partners, enabling modulation of both the specificity and the strength of the interaction. Coevolution could also explain how a JDP might have been able to switch Hsp70 partnership during evolution, such as occurred in yeast mitochondria - switching of Hsc20 from interacting with a general multifunctional Hsp70 (Ssc1) to a specialized Hsp70 (Ssq1)^40^. A JDP that interacts with a given Hsp70 partner could gain the ability to interact with a “new” Hsp70 (i.e. product of gene duplication), upon emergence of mutation(s) that facilitate a new interaction, without affecting the initial one, at least during the period of transition^41^.

What effect might this diversity have had on the functionality of Hsp70 chaperone systems more generally? We suggest that it has allowed evolution of complex Hsp70/JDP networks, both in situations in which a single type of Hsp70 interacts with many different JDPs and in which more than one Hsp70 coexist in a cellular compartment. In the first situation, replacement could well modulate the strength of a JDP/Hsp70 interaction, explaining why some JDPs have “easier access” to their partner Hsp70 than others. Here, electrostatic interactions could play a particularly important role. A JDP with a highly charged J-domain binding face would be able to recognize its Hsp70 partner at a longer distance, giving it priority over others in forming a productive interaction. In the second situation, the diversity serves to “insulate” Hsp70 networks, each Hsp70 interacting effectively only with its own subset of JDPs. For example, it could explain how yeast cytosolic Hsp70s Ssa and Ssb, which share common ancestry, evolved into independent networks, each with its own JDP partners^42,43^.

In the two examples studied here, the predominant J-domain interactions are via helix II. However, their binding interfaces differ - the specialized Hsc20/Ssq1 interface involving mostly electrostatic interactions and the DnaJ^JD^/DnaK interface being a mosaic of electrostatic and hydrophobic interactions ^22^. However, across JDP/Hsp70s partnerships the interfaces may be even more varied. Interestingly, in two cases, Helix III of the J-domain was found to be very important for interaction with its Hsp70 partner - auxilin^33,44^ and polyomavirus T Antigen^45^. As only a few J-domain/Hsp70 interfaces have been studied, the breath of sequence variability, that supports proper positioning of the HPD sequence, the key to functional partnership, may be larger still.

### Concluding remarks

Overall our data suggests a two-step process of J-domain function: initial binding that forms similar poses via highly variable residue-residue contacts, followed by a mechanistic step in which the invariant HPD perturbs the Hsp70 NBD-SBDβ,linker interface to foster a key allosteric transition. This separation could have future importance beyond a basic understanding of molecular chaperone function. It has long been appreciated that Hsp70 systems impinge on many biological functions and thus, not surprisingly, connected to many pathologic conditions raging from cancer, metabolic diseases to aging^46,47^. As it has become appreciated that many Hsp70s partner with multiple JDPs^1,43^, the idea of targeting a JDP, rather than Hsp70 itself, has emerged^48^. Our findings showing a clear separation between recognition and mechanistic function provide a possible avenue to approach such intervention – modulation of the recognition step without affecting the mechanistic step, thus, manipulating the specificity or strength of particular J-domain/Hsp70 interactions.

## Methods

### Molecular Dynamics (MD) Simulations

All simulations were performed using Gromacs 5^49^ with the Plumed 2.1 plugin^50^. If not stated otherwise, the CHARMM36 force field^51^ was used for proteins, ions and Mg-ATP, and the TIP3P model was used for water. In each of the simulation boxes, the numbers of Na^+^ and Cl^-^ ions were adjusted to 0.15 M. Temperature was kept at 310 K with the v-rescale algorithm^52^ using a coupling constant of 0.1 ps. Pressure was kept at 1 bar using the Parrinello-Rahman algorithm^53^ with a coupling time of 5 ps. Periodic boundary conditions were applied and the Particle Mesh Ewald summation^54^ was used to calculate long-range electrostatic interactions with a cut-off radius of 1 nm and a Fourier grid spacing of 0.12 nm. Van der Waals interactions were calculated with Lennard-Jones potential with a cut-off radius of 1 nm. All bonds involving hydrogen were constrained using the LINCS algorithm. Leap-frog Verlet algorithm was used to integrate equations of motion with a time step of 2 fs.

### Ssq1 homology model

Ssq1 model was built with I-TASSER^55^ using the ATP-bound structure of DnaK (PDB ID 4B9Q) as a template with C-score 0.34. The Mg-ATP complex and structural water molecules were placed in the conserved ATP binding pocket of Ssq1 according to the DnaK structure (PDB ID 4B9Q). The obtained model of Ssq1 was refined by 3-μs-long MD simulation with position restraints applied to Mg-ATP and structural water during the first 1 μs to ensure a proper relaxation of the ATP-binding pocket.

### Model of Hsc20-Ssq1 complex

The structures of Hsc20 (PDB ID 3UO3) and Ssq1 (homology model) used for molecular docking were first subjected to unbiased MD simulations to sample global protein structural dynamics. The Hsc20 simulation system was composed of a single protein in a 9.2 nm × 9.2 nm × 9.2 nm dodecahedron box filled with ∼17000 water molecules, while the Ssq1 simulation system was composed of a single ATP-bound protein solvated with ∼48,500 water molecules in a 13 nm × 13 nm × 13 nm dodecahedron box. Both systems were simulated using CHARMM36^51^ for 10 and 2 μs, respectively, and independently, using AMBER99SB-ILDN^56^ for 5 and 2 μs, respectively, to test for force-field dependence. The obtained MD trajectories were separately clustered with g_cluster using a 0.35 nm RMSD cut off for Hsc20 and a 0.7 nm RMSD cut off for Ssq1 (to account for large conformational fluctuations of the sub-domain SBDα). Centroids of each of the 12 and 7 clusters identified for Hsc20 and Ssq1, respectively, were treated as representative structures of the proteins and paired together in 84 independent docking runs using ClusPro^57^. As a result, 8,973 models were obtained. For each of the 84 docking runs, we selected the four best scoring models in each of the following criteria: a) binding energy estimation, b) minimal distance between the J-domain helix II and the NBD, c) minimal distance between the J-domain helix III and the NBD, d) minimal distance between the J-domain and the NBD. Redundancy among 336 selected models was removed with g_cluster, which resulted in 33 distinct models of the Hsc20-Ssq1 complex. Next, each of these models was placed in a 16 nm × 16 nm × 16 nm dodecahedron box and solvated with ∼85000 water molecules. The resulting 33 systems were energy-minimized and the protein side chains were relaxed with 50 ns-long MD simulations, during which the protein backbone atoms were restrained. Finally, the systems were simulated by unbiased MD for a minimum of 500 ns (total time for 33 simulated systems was 21 μs). The resulting trajectories were clustered based on heavy atoms of helices II and III of the Hsc20 J-domain after superimposing the Hsc20-Ssq1 complex structures using all Cα atoms of the conformationally rigid NBD domain. This was done using g_cluster with a single linkage method and a RMSD cutoff of 0.18 nm to obtain a fine characterization of the bound-state ensemble.

### MD simulations of DnaK and the DnaJ^JD^-DnaK structures

The systems were composed of either DnaK (PDB ID 4B9Q) or DnaK in complex with the J-domain of DnaJ (PDB ID 5NRO) placed in 13 nm × 13 nm × 13 nm dodecahedron box and solvated with approx. 52,000 water molecules. Crystal water molecules coordinated to Mg^2+^ ion in Mg-ATP were kept. Both systems were simulated for 1 μs and the final structures from these simulations were used as initial configurations for the subsequent free energy computations.

### Energy landscape of intramolecular contacts with the conserved R^NBD^ upon J-domain binding to Hsp70

The free energy landscapes describing the formation of intramolecular ion pairs by the critical R^NBD^ (R207 in Ssq1 and R167 in DnaK) were computed with well-tempered multiple walkers 2D-metadynamics^58^. In order to examine the effect of the J-domain binding on the R^NBD^ interactions, the simulation systems contained either Hsc20-Ssq1 or DnaJ^JD^-DnaK complexes or individual proteins, Ssq1 or DnaK, respectively, placed in a 11.8 nm **×** 11.8 nm **×** 11.8 nm dodecahedron box filled with ∼34,000 water molecules. To make the system significantly smaller and thus achieve better convergence of the free energy maps, the lid sub-domain, SBDα, was truncated by removing residues 570-657 of Ssq1 and 535-602 of DnaK. As two coordinates for 2D-metadynamics, we used the center of mass (COM) distances between the guanidine group of the R^NBD^ residue and the carboxylic groups of the partner aspartate residues at the SBDβ (D517 in Ssq1 or D481 in DnaK) and at the interdomain linker (D429 in Ssq1 or D481 in DnaK). Biasing potential was added every 10 ps with the bias factor of 15, a height of 0.1 kJ and a width of 0.04 nm. The sampled range of both coordinates was restricted to a 0.3--1.9 nm interval by applying one-sided harmonic potentials with a force constant of 3000 kJ nm^-2^. For both systems, 10 independent 1-μs-long walker-simulations were carried out. The final free energy maps were averaged over 15 individual maps computed in 25-ns intervals from the last 350 ns of the simulations to average out free energy fluctuations^59^.

The free energy profiles for the transition of the conserved arginine residue from intramolecular ion pairs with SBDβ (D517 for Ssq1, D481 for DnaK) and the linker (D429 for Ssq1, D481 for DnaK) to an intermolecular contact with the aspartate in the HPD (D50 for Hsc20, D35 for DnaJ) were computed using well-tempered multiple walkers metadynamics. The sampled coordinate was defined as the difference between the two distances: i) the COM distance between the arginine (R^NBD^) guanidine group and the carboxylic groups of the aspartate residues in the linker (D^Lk^) and in the SBDβ (D^SBD^) and ii) the COM distance between the R^NBD^ guanidine group and the carboxylic group of the D^HPD^ (see Fig. 2). To speed up convergence, both of these component distance coordinates were restricted to up to 1.5 nm by applying one-sided harmonic potentials with a force constant of 3000 kJ nm^-2^. The bias potential was incremented every 10 ps by adding Gaussian-shaped potentials with height and width of 0.06 kJ and of 0.04 nm, respectively, and a bias factor of 8. For both systems, 10 walkers of 400 ns each were simulated at the same time. The final free energy profiles were averaged over 8 individual profiles computed in 20-ns intervals from the last 140 ns to limit free energy fluctuations.

### Conformational transition of Hsp70 upon J-domain binding

The free energy profiles describing the compactness of the NBD, linker and SBDβ interface in Ssq1 and DnaK in the presence and absence of J-domain were computed with replica-exchange umbrella sampling (REUS). The simulations of the protein complexes (Hsc20-Ssq1 and DnaJ^JD^-DnaK) and free proteins (Ssq1 and DnaK) were performed in a 11.8 nm **×** 11.8 nm **×** 11.8 nm dodecahedron box filled with ∼36,000 water molecules. The reaction coordinate was defined as the COM distance between the Cα atoms of SBDβ (residues 434-541 in Ssq1 and 395-505 in DnaK) and the Cα atoms of the NBD region at the interface with SBDβ (residues 112-143 and 186-262 in Ssq1, residues 75-104 and 146-226 in DnaK). To accelerate the free energy convergence, the lid sub-domain (SBDα) was truncated (residues 542-657 in Ssq1 and 506-602 in DnaK). For all systems, the initial configurations for REUS simulations were alternately selected from two independent 200 ns steered-MD simulations. In these simulations, the distance between SBDβ and NBD was gradually increased using a moving one-sided harmonic potential with a force constant of 2500 kJ nm^-2^, applied to either the COM distance or the minimal distance between the Cα atoms of SBDβ and the NBD interfacial residues. The reaction coordinate in the 2.5-2.9 nm range was divided into 5 equally spaced windows in which the system was restrained using a harmonic potential with a spring constant of 2500 kJ nm^-2^. The same approach was used to study the effect of the R^NBD^->D substitutions (Ssq1(R207D) and DnaK(R167D)) on the relative arrangement of NBD and SBDβ. To perform these simulations, the R^NBD^ residue was replaced by aspartate in all initial structures. For each of the REUS windows at least 800 ns long simulation were performed. Exchanges between neighboring umbrella sampling windows were attempted every 2 ps. The first 100 ns of the trajectory was discarded and free energy profiles were computed using WHAM^60^ with the Monte Carlo bootstrap method to estimate uncertainties of the free energy. The number of contacts between the domains (NBD/SBDβ or NBD/linker) is defined as the number of non-hydrogen atoms within a cutoff distance of 0.5 nm. To compute the number of contacts as a function of the COM distances between the domains, REUS trajectories were reweighted using a factor of exp((*U*_*i*_(*r*) − *F*_*i*_)/*kT*), where *U*_*i*_(*r*) denotes the value of the biasing potential for a given frame and *F*_*i*_ denotes the free energy added to the i-th US window by the applied bias.

### Hsc20-Ssq1 interaction analysis

All inter-protein interactions were analyzed using a 10.5 μs MD trajectory of the Hsc20-Ssq1 complex representing a dominant bound state. (i) Ion pairs between Hsc20 and Ssq1 were identified using a cut-off of 0.5 nm for the distance between the centers of mass of the interacting charged groups^61^. (ii) Hydrogen bonds were identified using a standard geometric criterion: a 0.35 nm cut-off for donor-acceptor distance and a 30° cut-off for hydrogen-donor-acceptor angle. (iii) The hydrophobic contact surface area was estimated using the g_sas tool from the GROMACS package^49^. The hydrophobic surface buried in the Hsc20/Ssq1 interface was computed as the difference between the surface of hydrophobic residues exposed to water in the dissociated and bound state. (iv) Relative contributions to the Hsc20-Ssq1 binding free energy of individual residues were calculated with g_mmpbsa^62^ using a single trajectory approach, non-linear Poisson-Boltzmann solver and the CHARMM charges and atomic radii.

### Spontaneous binding of Hsc20 to Ssq1 and binding free energy

To study spontaneous binding between Hsc20 and Ssq1, the simulation dodecahedron box of size 16.2 nm × 16.2 nm × 16.2 nm containing single copies of Hsc20 and ATP-bound Ssq1 solvated with approx. 96500 water molecules was prepared. The proteins were initially placed such that the COM distance between positively charged residues of the Hsc20 J-domain helix II (R37, R38, R41) and negatively charged residues of Ssq1 NBD (D246, E248, D249, E253, D362, D364) was equal to 3 nm. For this system, five independent unbiased MD simulations were performed which were terminated after 60 ns as this time was sufficient for the native complex to spontaneously form in 2 out of 5 replicas. The free energy profiles for Ssq1 interaction with Hsc20 and with Hsc20(R37A,R41A) substitution variant were computed using replica-exchange umbrella sampling (REUS)^63^. As a reaction coordinate for REUS simulations, we used the COM distance between the respective binding sites corresponding to residues 34-44 of Hsc20 (Helix II of the J-domain) and residues 244-255 and 425-429 of Ssq1 (NBD). The harmonic potential was used to restrain the system along the reaction coordinate (spring constants and centers of the potential are summarized in Supplementary Table S1) and the initial configurations were taken from the spontaneous binding trajectories. The same initial configurations were used for the Hsc20(R37A,R41A) variant obtained by substituting R37 and R41 to alanine in all initial frames. For each of the REUS windows 400 ns long simulations were performed. Exchanges between neighboring windows were attempted every 2 ps and accepted by the Metropolis criterion. The free energy profiles were determined from the last 350 ns of thus obtained trajectories using the standard weighted histogram analysis method (WHAM)^60^ and the Monte Carlo bootstrap method to estimate uncertainties of the free energy differences.

### Evolutionary analyses

Protein sequences were retrieved from OMA orthology database ^64^: Hsc20 (OMAGroup:684646), mitochondrial Hsp70 SSC1(OMAGroup:546796), DnaJ (OMAGroup:571688), DnaK (OMAGroup:546796) cytosolic Hsp70s SSA1(OMAGroup:555679), SSA2(OMAGroup:557653), SSA3(OMAGroup:556877), SSA4(OMAGroup:557493), endoplasmic reticulum Hsp70s KAR2 (OMAGroup:555433). Sequences were aligned using Clustal Omega v1.2.2 with default parameters^65^. Alignment logos were generated using the WebLogo server^66^. Ancestral states of sites homologous to D517 of Ssq1 and D481 of DnaK were reconstructed in FastML using empirical Bayes method with Maximum Likelihood reconstruction of insertions and deletions^67^.

To infer Hsp70 phylogeny, a multiple sequence alignment was converted into a Hidden Markov Model^68^ using hmmbuild program from the HMMER package. Forward–backward algorithm was used to compute a posterior probability (pp) for each site representing the degree of confidence in each position (residue or gap) of the alignment for each sequence. Amino acid positions with pp <0.7 were removed from the multiple sequence alignment^69^. 1,000 maximum likelihood (ML) searches were performed using RAxML v8.2.10^70^ with 1000 rapid bootstrap replicates, under the LG model of amino acid substitution and GAMMA model of rate heterogeneity with four discrete rate categories and the estimate of proportion of invariable sites (LG + I + G)^71^, which was determined as the best-fit model by ProtTest v3.2 following Akaike Information Criterion^72^. Hsp70s tree topology was used for all calculations.

Maximum likelihood implementation of the Coev model^32^ was used to test coevolution between amino acid sequence positions homologous to those occupied by residues that were at a cutoff distance of 0.5 nm across the Hsc20/Ssq1 and DnaJ^JD^/DnaK binding interfaces. Coevolution analyses were based on the Hsp70 phylogeny, with Hsc20 orthologs paired with mitochondrial Hsp70 partners and DnaJ orthologs paired with DnaK partners. The fit of the Coev model in comparison to the ‘independent’ model was tested using Δ Akaike information criterion (ΔAIC = AIC_independent_ - AIC_Coev_). To establish the threshold for the ΔAIC value to be accepted as evidence for coevolution^32^, we simulated sequence alignment based on the same phylogenetic tree as the original data but using the ‘independent’ model of sequence evolution. We used evolver software from the PAML 4^73^ for this simulation. The 99^th^percentile of this expected ΔAIC distribution provided thresholds to consider the observed ΔAIC for Coev model to be accepted as coevolving with p<0.01 confidence level.

### Protein purification

Expression of Bpa-containing Hsc20 variants (Q46Bpa and M51Bpa) with a polyhistidine tag at the C-terminus was performed using the *E. coli* strain C41(DE3) carrying two plasmids: pET21d encoding Hsc20 Bpa variant and pSUPT BpF encoding an engineered tRNA synthetase and tRNA_CUA_ for p-benzoyl-phenylalanine (Bpa) incorporation. Cells were cultured in the M9 minimal medium. Protein expression was induced by adding isopropyl-1-thio-D-galactopyranoside (IPTG) at a final concentration of 0.2 mM, L-arabinose at a final concentration of 0.2% (v/v) and Bpa at a final concentration of 0.1 mM. Cells were harvested and lysed using a French press in buffer J1 (20 mM Tris-HCl, pH 8.0, 500 mM NaCl, 1 mM phenylmethanesulfonylfluoride (PMSF), 10% glycerol, 30 mM imidazole, pH 8.0). After a clarifying spin, the supernatant was precipitated with ammonium sulfate (0.35 mg/ml). After centrifugation, the pellet was resuspended in buffer J1 and dialyzed overnight against buffer J1. Next, the proteins were subjected to HisBind Resin (Novagen) chromatography. After sequential washing steps with buffers J1, J2 (20 mM Tris-HCl, pH 8.0, 1 M NaCl, 1 mM PMSF, 10% glycerol, 30 mM imidazole, pH 8.0, 1 mM ATP, 2 mM MgCl_2_) and J1, proteins were eluted with a linear 30–300 mM imidazole gradient in buffer J1. Fractions containing Hsc20 were collected, pooled and dialyzed against buffer J3 (40 mM potassium phosphate, pH 6.8; 10% (v/v) glycerol; 75 mM NaCl; 0.05% Triton X-100; 5 mM β-mercaptoethanol). After dialysis proteins were loaded on a P11 cellulose column (Spark Scientific Ltd), washed with a 150 mM NaCl solution in J3 buffer and eluted with a linear 150-800 mM NaCl gradient in J3 buffer. Fractions containing the purest Hsc20 were collected, pooled and dialyzed against the final buffer (20 mM Tris-HCl pH 8.0, 10% glycerol, 50 mM NaCl; 5 mM β-mercaptoethanol). Aliquots were stored at –70 °C.

All other Hsc20 variants with a polyhistidine tag at the C-terminus were purified according to ^24^, except *E. coli* strain C41(DE3) was used for expression. Ssq1 variants were purified as described^74^. Mge1 was purified as described^24^. Isu1 from *Chaetomium thermophilum* (Ius1_Ct_) was purified as described^25^

In all cases, protein concentrations, determined by using the Bradford (Bio-Rad) assay system with bovine serum albumin as a standard, are expressed as the concentration of monomers.

### ATPase Activity of Ssq1

The ATPase activity of Ssq1 variants defective in the Hsc20/Ssq1 interaction was measured as described by ^24^ with 0.5 μM Ssq1, 500 μM PVK-p peptide (LSLP**PVK**LHC) derived from the Isu1 sequence containing the PVK motif that interacts with substrate binding site of Ssq1^75^, 0.5 μM Mge1, and Hsc20 at the indicated concentrations in buffer A (40 mM Hepes–KOH, pH 7.5, 100 mM KCl, 1 mM dithiothreitol, 10 mM MgCl_2_, and 10% (v/v) glycerol). Reactions (15 μl) were initiated by the addition of ATP (2 μCi, DuPont NEG-003H, 3000 Ci/mmol) to a final concentration of 120 μM. Incubation was carried out at 25°C, and the reaction was terminated after 15 min by the addition of 100 μl of 1M perchloric acid and 1mM sodium phosphate. After addition of 400 μl of 20 mM ammonium molybdate and 400 μl of isopropyl acetate, samples were mixed and the phases were separated by a short centrifugation. An aliquot of the organic phase (150 μl), containing the radioactive orthophosphate-molybdate complex, was removed and radioactivity was determined by liquid scintillation counting. Control reactions lacking protein were included in all experiments. Values were plotted in GraphPadPrism. The ATPase activity corresponding to the maximal stimulation (Vm) of the wild-type Hsc20 was set to 1.

The ATPase activity of the Ssq1(R207D) variant was measured using an enzyme-coupled spectrophotometric assay^76^. Reaction mixtures of 0.5 ml contained 1 μM Ssq1 R207D or WT, 1 mM ATP, 0.265 mM NADH, 100 U/ml lactic dehydrogenase, 70 U/ml pyruvate kinase and 2.8 mM phosphoenolpyruvate in buffer (50 mM Hepes-KOH, pH 7.5, 150 mM KCl, 20 mM magnesium acetate, 10 mM dithiothreitol). If indicated, Mge1 was at 1 μM, Isu1_Ct_ at 3 μM and Hsc20 at 3 μM. Reactions were initiated by mixing ATP with proteins and then incubating at 23°C. NADH absorbance was measured at 340 nm for 500 seconds in JASCO V-660 spectrophotometer.

### Site specific photo-crosslinking

Reaction mixtures (20 μl) containing 20 μM Hsc20 Bpa-containing variants and 5 μM Ssq1 variants were prepared in the reaction buffer (40 mM Hepes–KOH, pH 7.5, 100 mM KCl, 1 mM dithiothreitol, 10 mM MgCl_2_, and 10% (v/v) glycerol). Reactions were initiated by the addition of ATP to a final concentration of 2 mM and incubated at 25°C for 15 min. Reactions were then exposed to 254 nm UV light (CL-1000 Ultraviolet Crosslinker) for 7 minutes. Control reactions were not exposed to UV light. Next, 5 μl of 4-fold concentrated Laemmli sample buffer (125 mM Tris-HCl, pH 6.8, 5% sodium dodecyl sulfate, 10% 2-mercaptoethanol, 20% (v/v) glycerol) was added to the reaction mixes. After 10 min incubation at 100°C, the total reaction volume was loaded onto SDS-PAGE and visualized by Coomassie blue stain. Every experiment was replicated three times.

### Identification of photo-crosslinking products using mass spectroscopy (MS)

Gel pieces were dried with acetonitrile and subjected to reduction with 10 mM dithiothreitol in 100mM NH_4_HCO_3_ for 30 min at 57°C. Cysteines were then alkylated with 0.5 M iodoacetamide in 100 mM NH_4_HCO_3_ (45 minutes in a dark room at room temperature) and proteins were digested overnight with 10 ng/μl trypsin (CAT NO V5280, Promega) in 25 mM NH_4_HCO_3_ ^77^. Digestion was stopped by adding trifluoroacetic acid at a final concentration of 0.1%. The mixture was centrifuged at 4°C, 14 000 g for 30 min, to remove any remaining solid. MS analysis was performed by LC-MS in the Laboratory of Mass Spectrometry (Institute of Biochemistry and Biophysics, Polish Academy of Sciences, Warsaw) using a nanoAcquity UPLC system (Waters) coupled to an Orbitrap Elite (Thermo Fisher Scientific). The mass spectrometer was operated in the data-dependent MS2 mode, and data were acquired in the m/z range of 300-2000. Peptides were separated by a 180 min linear gradient of 95% solution A (0.1% formic acid in water) to 35% solution B (acetonitrile and 0.1% formic acid). The measurement of each sample was preceded by three washing runs to avoid cross-contamination. Data were searched with Mascot ^78^, with the following parameters: enzyme: Trypsin, parent ions mass tolerance: 20 ppm, fragment ion mass tolerance: 0.1, missed cleavages: 1, fixed modifications: Carbamidomethyl (C), variable modifications: Oxidation (M). Protein identification was validated using a target-decoy search strategy.

## Acknowledgements

We thank Xavier Meyer and Daniele Silvestro, University of Lausanne, for discussions about detecting coevolution. We thank Aneta Grabinska for technical help with the ATPase analyses.

This work was supported by POIR.04.04.00-00-4114 /17-00 project carried out within the TEAM programme co-financed by the European Union under the European Regional Development Fund (JM) and by the National Institutes of Health R35GM127009 (EAC). This research was supported in part by PL-Grid Infrastructure (http://www.plgrid.pl/en).

## Notes

#### Summary of Updates

author contact data and license updated

## References

(1) Kampinga, H. H.; Craig, E. A. The HSP70 chaperone machinery: J proteins as drivers of functional specificity. Nature Reviews Molecular Cell Biology 2010, 11 (8), 579.

(2) Rosenzweig, R.; Nillegoda, N. B.; Mayer, M. P.; Bukau, B. The Hsp70 chaperone network. Nature Reviews Molecular Cell Biology 2019, 20, 665.

(3) Clerico, E. M.; Meng, W.; Pozhidaeva, A.; Bhasne, K.; Petridis, C.; Gierasch, L. M. Hsp70 molecular chaperones: multifunctional allosteric holding and unfolding machines. Biochemical Journal 2019, 476 (11), 1653.

(4) Craig, E. A.; Marszalek, J. How Do J-Proteins Get Hsp70 to Do So Many Different Things? Trends in Biochemical Sciences 2017, 42 (5), 355.

(5) Kityk, R.; Kopp, J.; Sinning, I.; Mayer, M. P. Structure and Dynamics of the ATP-Bound Open Conformation of Hsp70 Chaperones. Molecular Cell 2012, 48 (6), 1.

(6) Qi, R.; Sarbeng, E. B.; Liu, Q.; Le, K. Q.; Xu, X.; Xu, H.; Yang, J.; Wong, J. L.; Vorvis, C.; Hendrickson, W. A. et al. Allosteric opening of the polypeptide-binding site when an Hsp70 binds ATP. Nature Structural & Molecular Biology 2013, 20 (7), 1.

(7) Mapa, K.; Sikor, M.; Kudryavtsev, V.; Waegemann, K.; Kalinin, S.; Seidel, C. A. M.; Neupert, W.; Lamb, D. C.; Mokranjac, D. The conformational dynamics of the mitochondrial Hsp70 chaperone. Molecular Cell 2010, 38 (1), 89.

(8) Marcinowski, M.; Höller, M.; Feige, M. J.; Baerend, D.; Lamb, D. C.; Buchner, J. Substrate discrimination of the chaperone BiP by autonomous and cochaperone-regulated conformational transitions. Nature Structural & Molecular Biology 2011, 18 (2), 150.

(9) Banerjee, R.; Jayaraj, G. G.; Peter, J. J.; Kumar, V.; Mapa, K. Monitoring conformational heterogeneity of the lid of DnaK substrate-binding domain during its chaperone cycle. The FEBS Journal 2016, 283 (15), 2853.

(10) Zhuravleva, A.; Clerico, E. M.; Gierasch, L. M. An interdomain energetic tug-of-war creates the allosterically active state in Hsp70 molecular chaperones. Cell 2012, 151 (6), 1296.

(11) Kityk, R.; Vogel, M.; Schlecht, R.; Bukau, B.; Mayer, M. P. Pathways of allosteric regulation in Hsp70 chaperones. Nature Communications 2015, 6, 8308.

(12) Vogel, M.; Bukau, B.; Mayer, M. P. Allosteric Regulation of Hsp70 Chaperones by a Proline Switch. Molecular Cell 2006, 21 (3), 359.

(13) General, I. J.; Liu, Y.; Blackburn, M. E.; Mao, W.; Gierasch, L. M.; Bahar, I. ATPase Subdomain IA Is a Mediator of Interdomain Allostery in Hsp70 Molecular Chaperones. PLoS Computational Biology 2014, 10 (5), e1003624.

(14) Vogel, M.; Mayer, M. P.; Bukau, B. Allosteric regulation of Hsp70 chaperones involves a conserved interdomain linker. Journal of Biological Chemistry 2006, 281 (50), 38705.

(15) Swain, J. F.; Dinler, G.; Sivendran, R.; Montgomery, D. L.; Stotz, M.; Gierasch, L. M. Hsp70 Chaperone Ligands Control Domain Association via an Allosteric Mechanism Mediated by the Interdomain Linker. Molecular Cell 2007, 26 (1), 27.

(16) Alderson, T. R.; Kim, J. H.; Cai, K.; Frederick, R. O.; Tonelli, M.; Markley, J. L. The Specialized Hsp70 (HscA) Interdomain Linker Binds to Its Nucleotide-Binding Domain and Stimulates ATP Hydrolysis in Both cisand transConfigurations. Biochemistry 2014, 53 (46), 7148.

(17) English, C. A.; Sherman, W.; Meng, W.; Gierasch, L. M. The Hsp70 interdomain linker is a dynamic switch that enables allosteric communication between two structured domains. Journal of Biological Chemistry 2017, 292 (36), 14765.

(18) Lai, A. L.; Clerico, E. M.; Blackburn, M. E.; Patel, N. A.; Robinson, C. V.; Borbat, P. P.; Freed, J. H.; Gierasch, L. M. Key features of an Hsp70 chaperone allosteric landscape revealed by ion-mobility native mass spectrometry and double electron-electron resonance. Journal of Biological Chemistry 2017, 292 (21), 8773.

(19) Liberek, K.; Marszalek, J.; Ang, D.; Georgopoulos, C.; Zylicz, M. Escherichia coli DnaJ and GrpE heat shock proteins jointly stimulate ATPase activity of DnaK. Proceedings of the National Academy of Sciences of the United States of America 1991, 2874.

(20) Suh, W. C.; Burkholder, W. F.; Lu, C. Z.; Zhao, X.; Gottesman, M. E.; Gross, C. A. Interaction of the Hsp70 molecular chaperone, DnaK, with its cochaperone DnaJ. Proceedings of the National Academy of Sciences of the United States of America 1998, 95 (26), 15223.

(21) Gassler, C. S.; Buchberger, A.; Laufen, T.; Mayer, M. P.; Schröder, H.; Valencia, A.; Bukau, B. Mutations in the DnaK chaperone affecting interaction with the DnaJ cochaperone. Proceedings of the National Academy of Sciences of the United States of America 1998, 95 (26), 15229.

(22) Kityk, R.; Kopp, J.; Mayer, M. P. Molecular Mechanism of J-Domain-Triggered ATP Hydrolysis by Hsp70 Chaperones. Molecular Cell 2018, 69 (2), 227.

(23) Malinverni, D.; Jost Lopez, A.; De Los Rios, P.; Hummer, G.; Barducci, A. Modeling Hsp70/Hsp40 interaction by multi-scale molecular simulations and coevolutionary sequence analysis. eLife 2017, 6, 19.

(24) Dutkiewicz, R.; Schilke, B.; Knieszner, H.; Walter, W.; Craig, E. A.; Marszalek, J. Ssq1, a mitochondrial Hsp70 involved in iron-sulfur (Fe/S) center biogenesis. Similarities to and differences from its bacterial counterpart. Journal of Biological Chemistry 2003, 278 (32), 29719.

(25) Delewski, W.; Paterkiewicz, B.; Manicki, M.; Schilke, B.; Tomiczek, B.; Ciesielski, S. J.; Nierzwicki, L.; Czub, J.; Dutkiewicz, R.; Craig, E. A. et al. Iron–Sulfur Cluster Biogenesis Chaperones: Evidence for Emergence of Mutational Robustness of a Highly Specific Protein–Protein Interaction. Molecular Biology and Evolution 2016, 33 (3), 643.

(26) Dror, R. O.; Dirks, R. M.; Grossman, J. P.; Xu, H.; Shaw, D. E. Biomolecular Simulation: A Computational Microscope for Molecular Biology. Annual Review of Biophysics 2012, 41 (1), 429.

(27) Barducci, A.; Bonomi, M.; Parrinello, M. Metadynamics. Wiley Online Library 2011, DOI: 10.1002/wcms.31 10.1002/wcms.31.

(28) Valsson, O.; Tiwary, P.; Parrinello, M. Enhancing Important Fluctuations: Rare Events and Metadynamics from a Conceptual Viewpoint. Annual Review of Physical Chemistry 2016, 67 (1), 159.

(29) Donald, J. E.; Kulp, D. W.; DeGrado, W. F. Salt bridges: geometrically specific, designable interactions. Proteins: Structure, Function, and Bioinformatics 2011, 79 (3), 898.

(30) Krishnamurthy, M.; Dugan, A.; Nwokoye, A.; Fung, Y.-H.; Lancia, J. K.; Majmudar, C. Y.; Mapp, A. K. Caught in the act: covalent cross-linking captures activator-coactivator interactions in vivo. ACS Chemical Biology 2011, 6 (12), 1321.

(31) de Juan, D.; Pazos, F.; Valencia, A. Emerging methods in protein co-evolution. Nature Reviews Genetics 2013, 14 (4), 1.

(32) Dib, L.; Silvestro, D.; Salamin, N. Evolutionary footprint of coevolving positions in genes. Bioinformatics 2014, 30 (9), 1241.

(33) Jiang, J.; Maes, E. G.; Taylor, A. B.; Wang, L.; Hinck, A. P.; Lafer, E. M.; Sousa, R. Structural Basis of J Cochaperone Binding and Regulation of Hsp70. Molecular Cell 2007, 28 (3), 422.

(34) Zhuravleva, A.; Gierasch, L. M. Substrate-binding domain conformational dynamics mediate Hsp70 allostery. Proceedings of the National Academy of Sciences of the United States of America 2015, 112 (22), E2865.

(35) Sikor, M.; Mapa, K.; von Voithenberg, L. V.; Mokranjac, D.; Lamb, D. C. Real-time observation of the conformational dynamics of mitochondrial Hsp70 by spFRET. The EMBO Journal 2013, 32 (11), 1639.

(36) Wieteska, L.; Shahidi, S.; Zhuravleva, A. Allosteric fine-tuning of the conformational equilibrium poises the chaperone BiP for post-translational regulation. eLife 2017, 6, 18966.

(37) Meng, W.; Clerico, E. M.; McArthur, N.; Gierasch, L. M. Allosteric landscapes of eukaryotic cytoplasmic Hsp70s are shaped by evolutionary tuning of key interfaces. Proceedings of the National Academy of Sciences of the United States of America 2018, 115 (47), 11970.

(38) Stetz, G.; Verkhivker, G. M. Computational Analysis of Residue Interaction Networks and Coevolutionary Relationships in the Hsp70 Chaperones: A Community-Hopping Model of Allosteric Regulation and Communication. PLoS Computational Biology 2017, 13 (1), e1005299.

(39) Bascos, N. A. D.; Mayer, M. P.; Bukau, B.; Landry, S. J. The Hsp40 J-domain modulates Hsp70 conformation and ATPase activity with a semi-elliptical spring. Protein Science 2017, 26 (9), 1838.

(40) Schilke, B.; Williams, B.; Knieszner, H.; Pukszta, S.; D’Silva, P.; Craig, E. A.; Marszalek, J. Evolution of Mitochondrial Chaperones Utilized in Fe-S Cluster Biogenesis. Current Biology 2006, 16 (16), 1660.

(41) Pukszta, S.; Schilke, B.; Dutkiewicz, R.; Kominek, J.; Moczulska, K.; Stepien, B.; Reitenga, K. G.; Bujnicki, J. M.; Williams, B.; Craig, E. A. et al. Co-evolution-driven switch of J-protein specificity towards an Hsp70 partner. EMBO Reports 2010, 11 (5), 360.

(42) Meyer, A. E.; Hung, N.-J.; Yang, P.; Johnson, A. W.; Craig, E. A. The specialized cytosolic J-protein, Jjj1, functions in 60S ribosomal subunit biogenesis. Proceedings of the National Academy of Sciences of the United States of America 2007, 104 (5), 1558.

(43) Sahi, C.; Craig, E. A. Network of general and specialty J protein chaperones of the yeast cytosol. Proceedings of the National Academy of Sciences of the United States of America 2007, 104 (17), 7163.

(44) Sousa, R.; Jiang, J.; Lafer, E. M.; Lafer, E. M.; Hinck, A. P.; Hinck, A. P.; Wang, L.; Taylor, A. B.; Taylor, A. B.; Maes, E. G. et al. Evaluation of competing J domain:Hsp70 complex models in light of existing mutational and NMR data. Proceedings of the National Academy of Sciences of the United States of America 2012, 109 (13), E734.

(45) Garimella, R.; Liu, X.; Qiao, W.; Liang, X.; Zuiderweg, E. R. P.; Riley, M. I.; Van Doren, S. R. Hsc70 Contacts Helix III of the J Domain from Polyomavirus T Antigens: Addressing a Dilemma in the Chaperone Hypothesis of How They Release E2F from pRb †. Biochemistry 2006, 45 (22), 6917.

(46) Zarouchlioti, C.; Parfitt, D. A.; Li, W.; Gittings, L. M.; Cheetham, M. E. DNAJ Proteins in neurodegeneration: essential and protective factors. Philosophical Transactions of the Royal Society of London. Series B, Biological Sciences 2017, 373 (1738), 20160534.

(47) Calderwood, S. K.; Gong, J. Heat Shock Proteins Promote Cancer: It’s a Protection Racket. Trends in Biochemical Sciences 2016, 41 (4), 311.

(48) Li, X.; Shao, H.; Taylor, I. R.; Gestwicki, J. E. Targeting Allosteric Control Mechanisms in Heat Shock Protein 70 (Hsp70). Current Topics in Medicinal Chemistry 2016, 16 (25), 2729.

(49) Abraham, M. J.; Murtola, T.; Schulz, R.; Páll, S.; Smith, J. C.; Hess, B.; Lindahl, E. GROMACS: High performance molecular simulations through multi-level parallelism from laptops to supercomputers. SoftwareX 2015, 1-2, 19.

(50) Bonomi, M.; Branduardi, D.; Bussi, G.; Camilloni, C.; Provasi, D.; Raiteri, P.; Donadio, D.; Marinelli, F.; Pietrucci, F.; Broglia, R. A. et al. PLUMED: a portable plugin for free-energy calculations with molecular dynamics Computer Physics Communication 2009, 180 (10), 1961.

(51) Huang, J.; MacKerell Jr, A. D. CHARMM36 all-atom additive protein force field: Validation based on comparison to NMR data. Journal of Computational Chemistry 2013, 34 (25), 2135.

(52) Bussi, G.; Donadio, D.; Parrinello, M. Canonical sampling through velocity rescaling. The Journal of Chemical Physics 2007, 126 (1), 014101.

(53) Parrinello, M.; Rahman, A. Polymorphic transitions in single crystals: A new molecular dynamics method. Journal of Applied Physics 1981, 52 (12), 7182.

(54) Darden, T.; York, D.; Pedersen, L. Particle mesh Ewald: An N·log(N) method for Ewald sums in large systems. The Journal of Chemical Physics 1993, 98 (12), 10089.

(55) Yang, J.; Zhang, Y. Protein Structure and Function Prediction Using I-TASSER. Current Protocols in Bioinformatics 2015, 52, 5.8.1.

(56) Lindorff-Larsen, K.; Piana, S.; Palmo, K.; Maragakis, P.; Klepeis, J. L.; Dror, R. O.; Shaw, D. E. Improved side-chain torsion potentials for the Amber ff99SB protein force field. Proteins: Structure, Function, and Bioinformatics 2010, 78 (8), 1950.

(57) Kozakov, D.; Hall, D. R.; Xia, B.; Porter, K. A.; Padhorny, D.; Yueh, C.; Beglov, D.; Vajda, S. The ClusPro web server for protein-protein docking. Nature Protocols 2017, 12 (2), 255.

(58) Raiteri, P.; Laio, A.; Gervasio, F. L.; Micheletti, C.; Parrinello, M. Efficient Reconstruction of Complex Free Energy Landscapes by Multiple Walkers Metadynamics †. The Journal of Physical Chemistry B 2006, 110 (8), 3533.

(59) Laio, A.; Gervasio, F. L. Metadynamics: a method to simulate rare events and reconstruct the free energy in biophysics, chemistry and material science. Reports on Progress in Physics 2008, 71 (12), 126601.

(60) Kumar, S.; Bouzida, D.; Swendsen, R. H.; Kollman, P. A.; Rosenberg, J. M. The Weighted Histogram Analysis Method for Free-Energy Calculations on Biomolecules. Journal of Computational Chemistry 1992, 13, 1011.

(61) Kumar, S.; Nussinov, R. Relationship between ion pair geometries and electrostatic strengths in proteins. Biophysical Journal 2002, 83 (3), 1595.

(62) Kumari, R.; Kumar, R.; Consortium, O. S. D. D.; Lynn, A. g_mmpbsa—A GROMACS Tool for High-Throughput MM-PBSA Calculations. Journal of Chemical Information and Modeling 2014, 54 (7), 1951.

(63) Sugita, Y.; Kitao, A.; Okamoto, Y. Multidimensional replica-exchange method for free-energy calculations. Journal of Chemical Physics 2000, 113 (15), 6042.

(64) Altenhoff, A. M.; Glover, N. M.; Train, C.-M.; Kaleb, K.; Warwick Vesztrocy, A.; Dylus, D.; de Farias, T. M.; Zile, K.; Stevenson, C.; Long, J. et al. The OMA orthology database in 2018: retrieving evolutionary relationships among all domains of life through richer web and programmatic interfaces. Nucleic Acids Research 2018, 46 (D1), D477.

(65) Sievers, F.; Wilm, A.; Dineen, D.; Gibson, T. J.; Karplus, K.; Li, W.; Lopez, R.; McWilliam, H.; Remmert, M.; ding, J. S. o. et al. Fast, scalable generation of high-quality protein multiple sequence alignments using Clustal Omega. Molecular Systems Biology 2011, 7, 1.

(66) Crooks, G. E.; Hon, G.; Chandonia, J.-M.; Brenner, S. E. WebLogo: a sequence logo generator. Genome Research 2004, 14 (6), 1188.

(67) Ashkenazy, H.; Penn, O.; Doron-Faigenboim, A.; Cohen, O.; Cannarozzi, G.; Zomer, O.; Pupko, T. FastML: a web server for probabilistic reconstruction of ancestral sequences. Nucleic Acids Research 2012, 40, W580.

(68) Eddy, S. R. Profile hidden Markov models Bioinformatics 1998, 14 (9), 755.

(69) Sarangi, G. K.; Romagné, F.; Castellano, S. Distinct Patterns of Selection in Selenium-Dependent Genes between Land and Aquatic Vertebrates. Molecular Biology and Evolution 2018, 35 (7), 1744.

(70) Stamatakis, A. RAxML version 8: a tool for phylogenetic analysis and post-analysis of large phylogenies. Bioinformatics 2014, 30 (9), 1312.

(71) Le, S. Q.; Gascuel, O. An Improved General Amino Acid Replacement Matrix. Molecular Biology and Evolution 2008, 25 (7), 1307.

(72) Darriba, D.; Taboada, G. L.; Doallo, R.; Posada, D. ProtTest 3: fast selection of best-fit models of protein evolution. Bioinformatics 2011, 27 (8), 1164.

(73) Yang, Z. PAML 4: Phylogenetic Analysis by Maximum Likelihood. Molecular Biology and Evolution 2007, 24 (8), 1586.

(74) Manicki, M.; Majewska, J.; Ciesielski, S.; Schilke, B.; Blenska, A.; Kominek, J.; Marszalek, J.; Craig, E. A.; Dutkiewicz, R. Overlapping Binding Sites of the Frataxin Homologue Assembly Factor and the Heat Shock Protein 70 Transfer Factor on the Isu Iron-Sulfur Cluster Scaffold Protein. Journal of Biological Chemistry 2014, 289 (44), 30268.

(75) Knieszner, H.; Schilke, B.; Dutkiewicz, R.; D’Silva, P.; Cheng, S.; Ohlson, M.; Craig, E. A.; Marszalek, J. Compensation for a defective interaction of the hsp70 ssq1 with the mitochondrial Fe-S cluster scaffold isu. Journal of Biological Chemistry 2005, 280 (32), 28966.

(76) Nørby, J. G. Coupled assay of Na+,K+-ATPase activity. Methods in Enzymology 1988, 156, 116.

(77) Orlowska, K. P.; Klosowska, K.; Szczesny, R. J.; Cysewski, D.; Krawczyk, P. S.; Dziembowski, A. A new strategy for gene targeting and functional proteomics using the DT40 cell line. Nucleic Acids Research 2013, 41 (17), e167.

(78) Perkins, D. N.; Pappin, D. J.; Creasy, D. M.; Cottrell, J. S. Probability-based protein identification by searching sequence databases using mass spectrometry data. Electrophoresis 1999, 20 (18), 3551.

