## Supplementary Data for "Two-step mechanism of J-domain action in driving Hsp70 function"

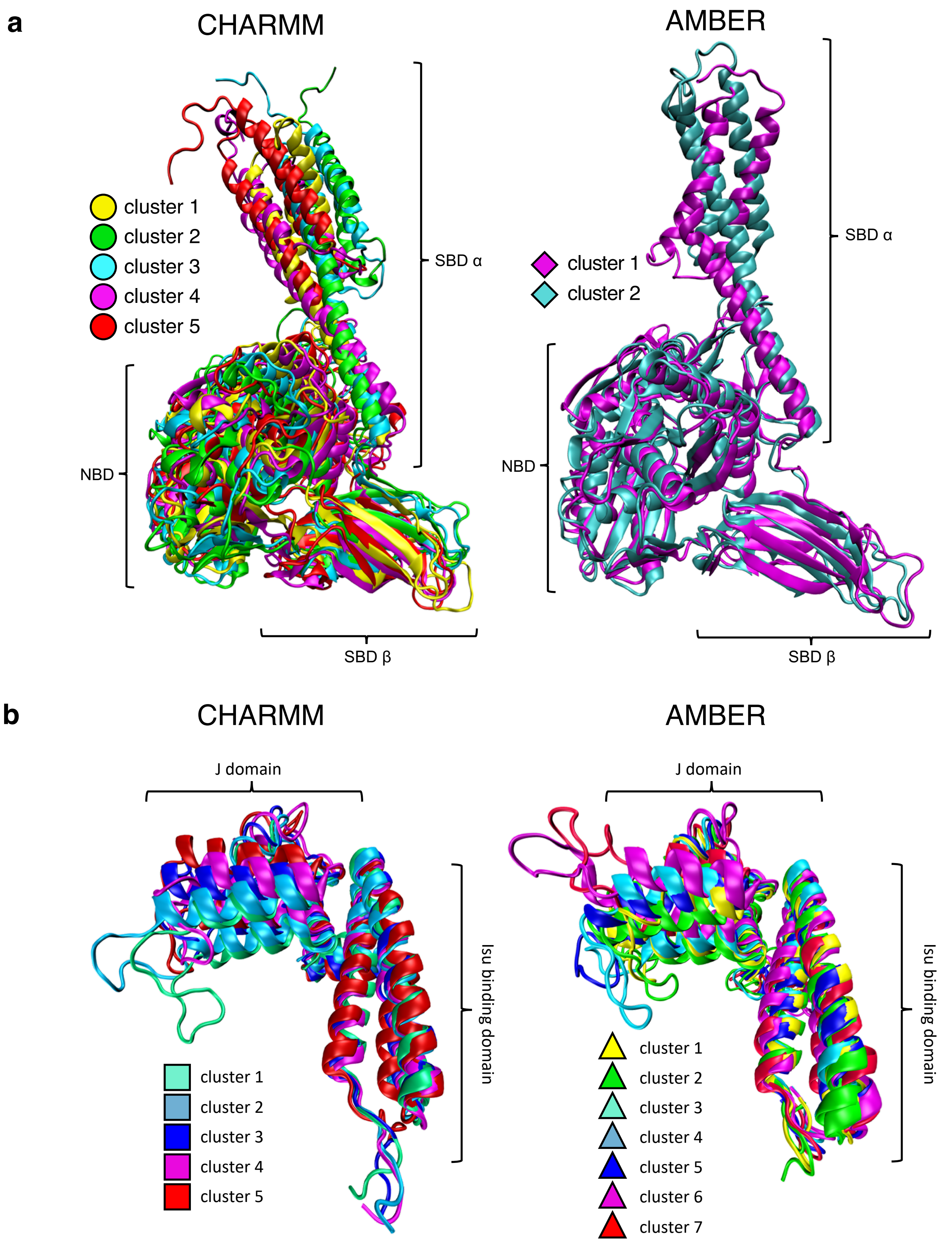


**Supplementary Fig. S1**

Representative structures of Ssq1 (a) and Hsc20 (b) from our unbiased molecular dynamics simulations carried out using the CHARMM36 (left) and AMBER99SB-ILDN (right) force fields. Ssq1 and Hsc20 were simulated for 2 and 10 μs, respectively, with CHARMM, and for 2 and 5 μs, respectively, with AMBER. The obtained trajectories were subject to cluster analysis with an RMSD cutoff of 0.7 nm (Ssq1) and 0.35 nm (Hsc20). The centroids of each cluster were superimposed and are shown in different colors. NBD, SBD$\alpha$ and SBD$\beta$ domains (in Ssq1) and J-domain and Isu1 binding domain (in Hsc20) are indicated.

**a**

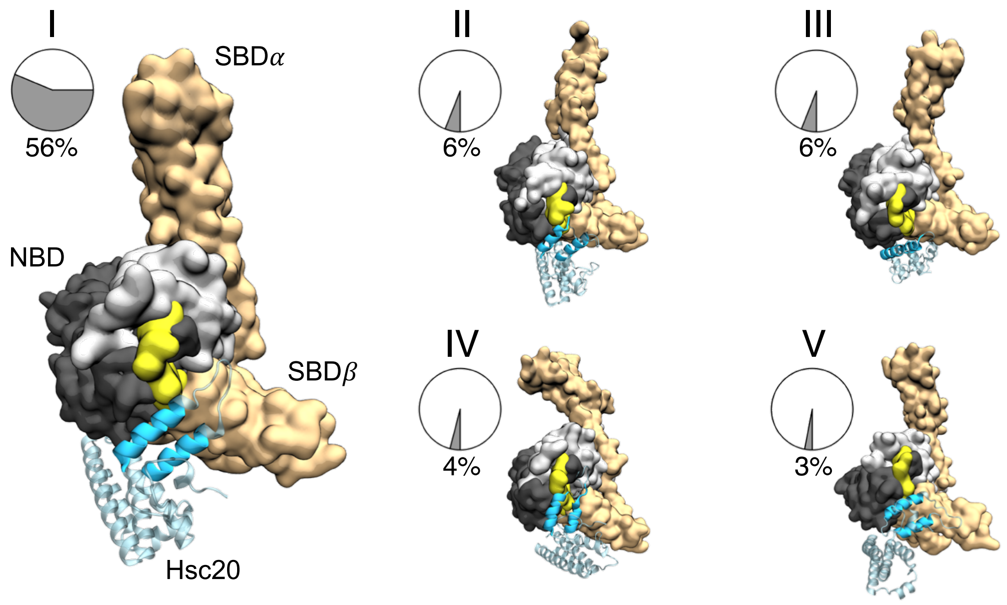


**b**

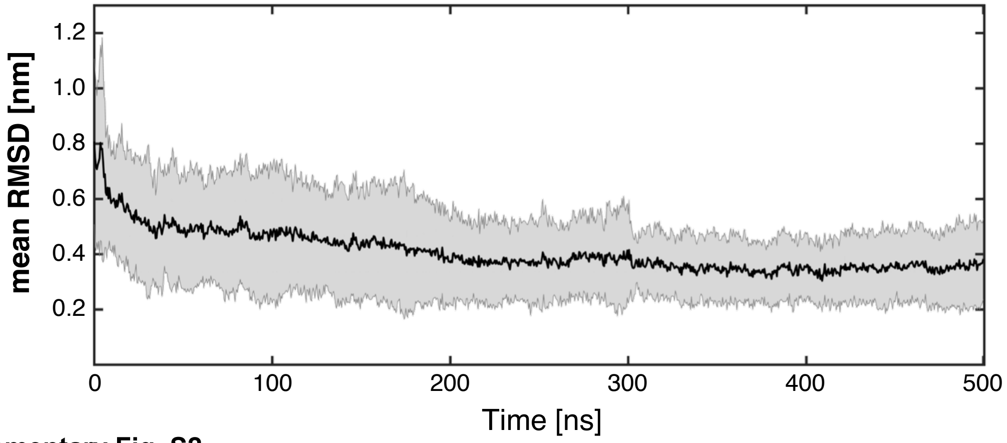


**Supplementary Fig. S2**

(a) Five most populated binding poses of Hsc20 along with their percentage contributions to the bound-state ensemble of the Hsc20-Ssq1 complex, obtained by the combined docking/MD simulation approach. Helices II and III are shown in cyan and the rest of Hsc20 in transparent cyan. SBD is in light brown, Ia and IIa subdomains of NBD are in light and dark grey, respectively, linker in yellow.

(b) Time evolution of the average RMSD of helices II and III of the J-domain with respect to the center of the most populated binding pose above (the dominant bound state). Averaging was done over all 18 independent MD trajectories that were found to visit the dominant bound state at least once over the course of 500 ns. The shaded area shows the standard deviation of the RMSD.

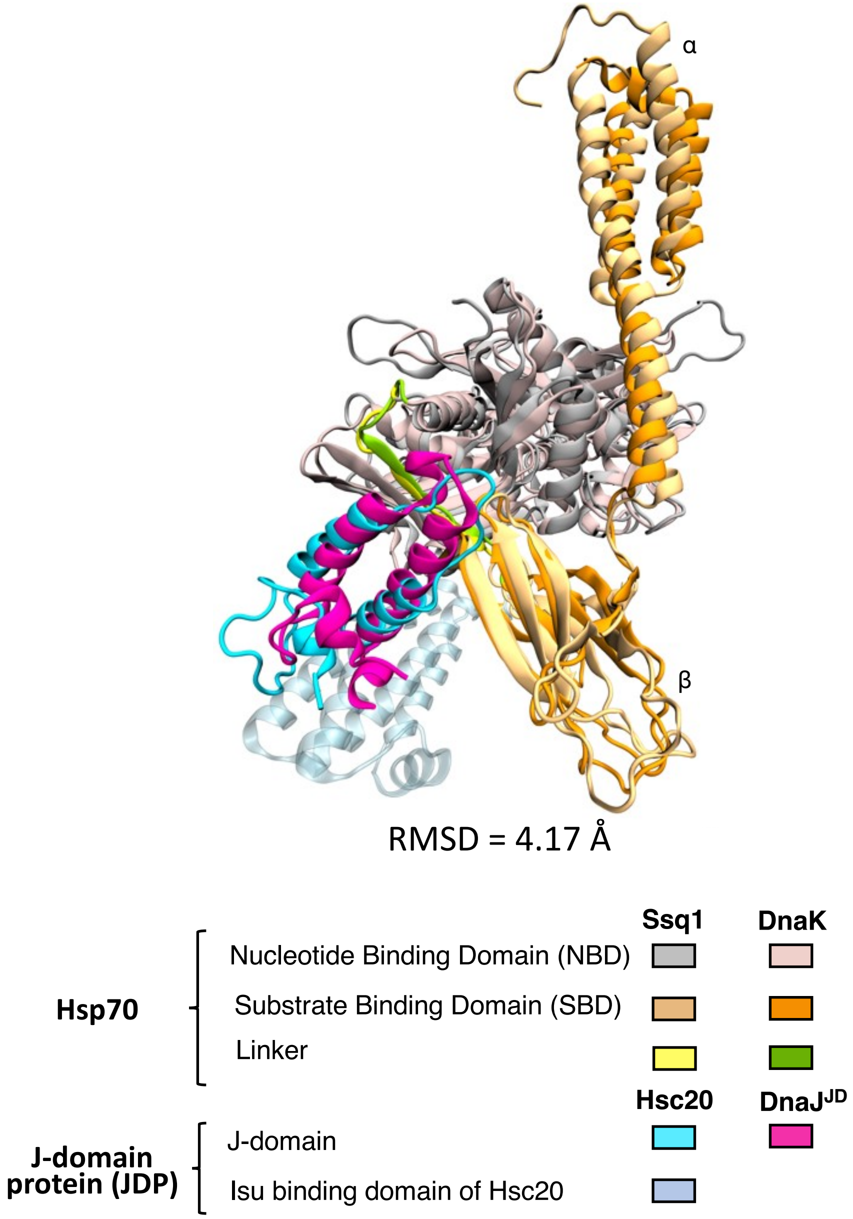


**Supplementary Fig. S3**

Structural alignment of the predicted Hsc20-Ssq1 complex and DnaJ$^{JD}$-DnaK crystal structure (the backbone RMSD of helices II and III of the J-domains after aligning NBDs is 4.17 Å). The J-domain, NBD, linker and SBD of aligned proteins are represented in cyan/magenta, grey/pink, yellow/green and light/dark orange for Hsc20-Ssq1 and DnaJ$^{JD}$-DnaK, respectively.

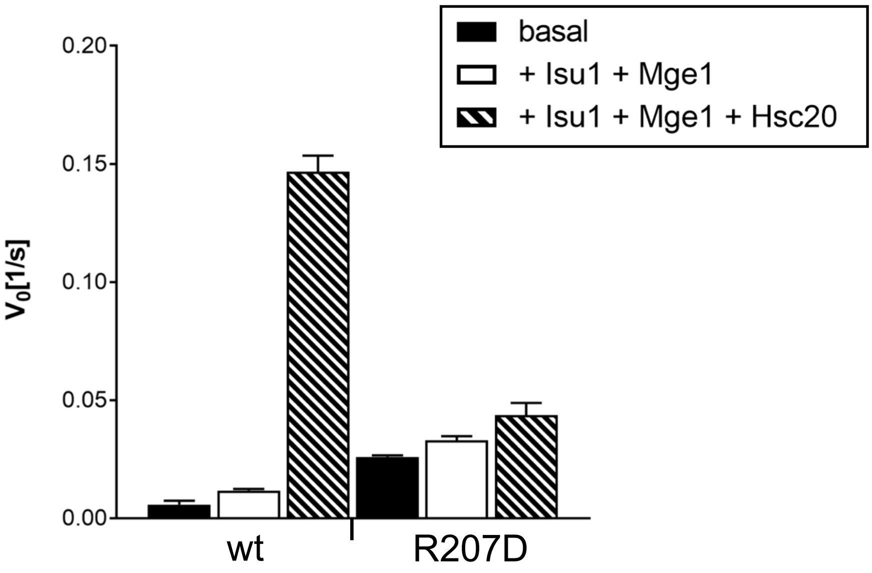


**Supplementary Fig. S4**
ATPase activity of Ssq1(R207D) variant. Steady-state ATPase activity of Ssq1 was measured alone, or in the presence of indicated proteins, using an enzymatically coupled assay. Bars represent average value for three independent measurements with error bars as standard deviation. Concentration of components: 1 μM Ssq1; 3 μM Hsc20; 3 μM Isu1; 1 μM Mge1; 1 mM ATP.

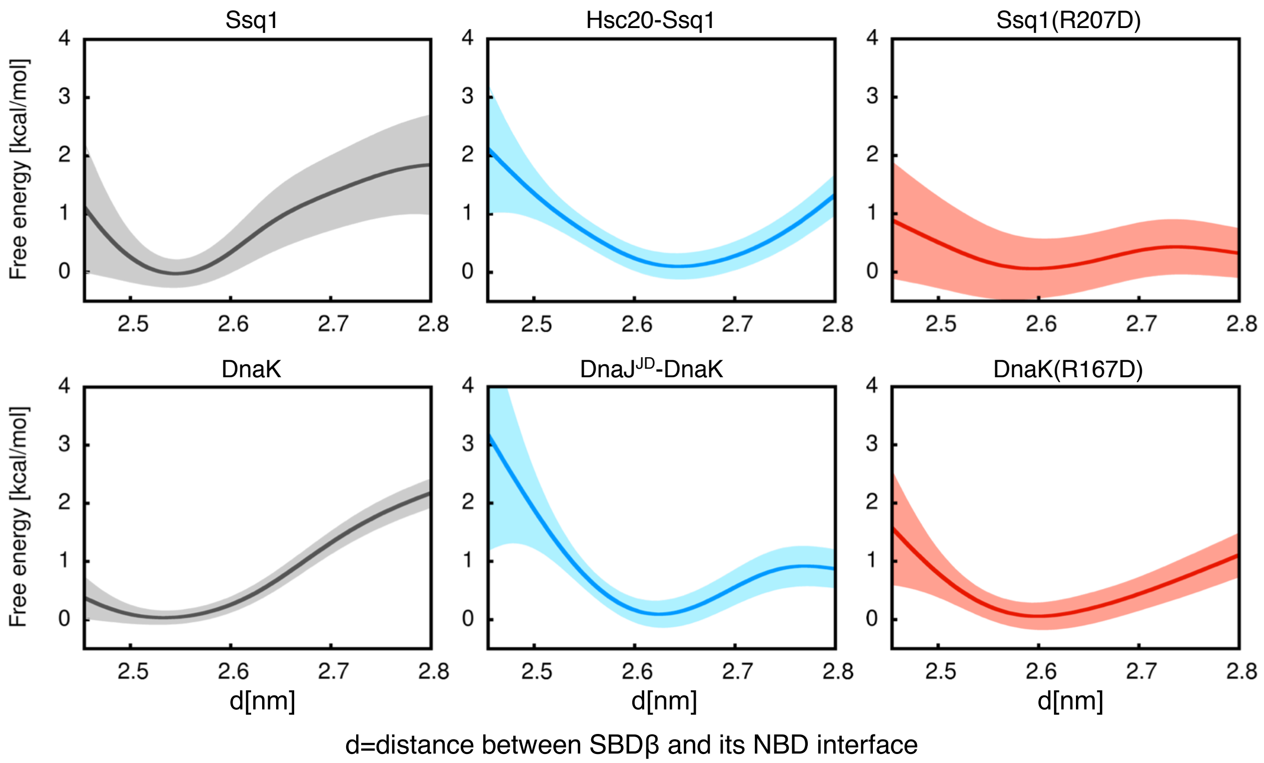


**Supplementary Fig. S5**

Free energy profiles as a function of the distance *d* (defined in Fig. 3) for Ssq1 and DnaK in red, for Hsc20-Ssq1 and DnaJ$^{JD}$-DnaK complexes in blue, and for Ssq1 and DnaK R$^{NBD}$->D substitution variants in purple.

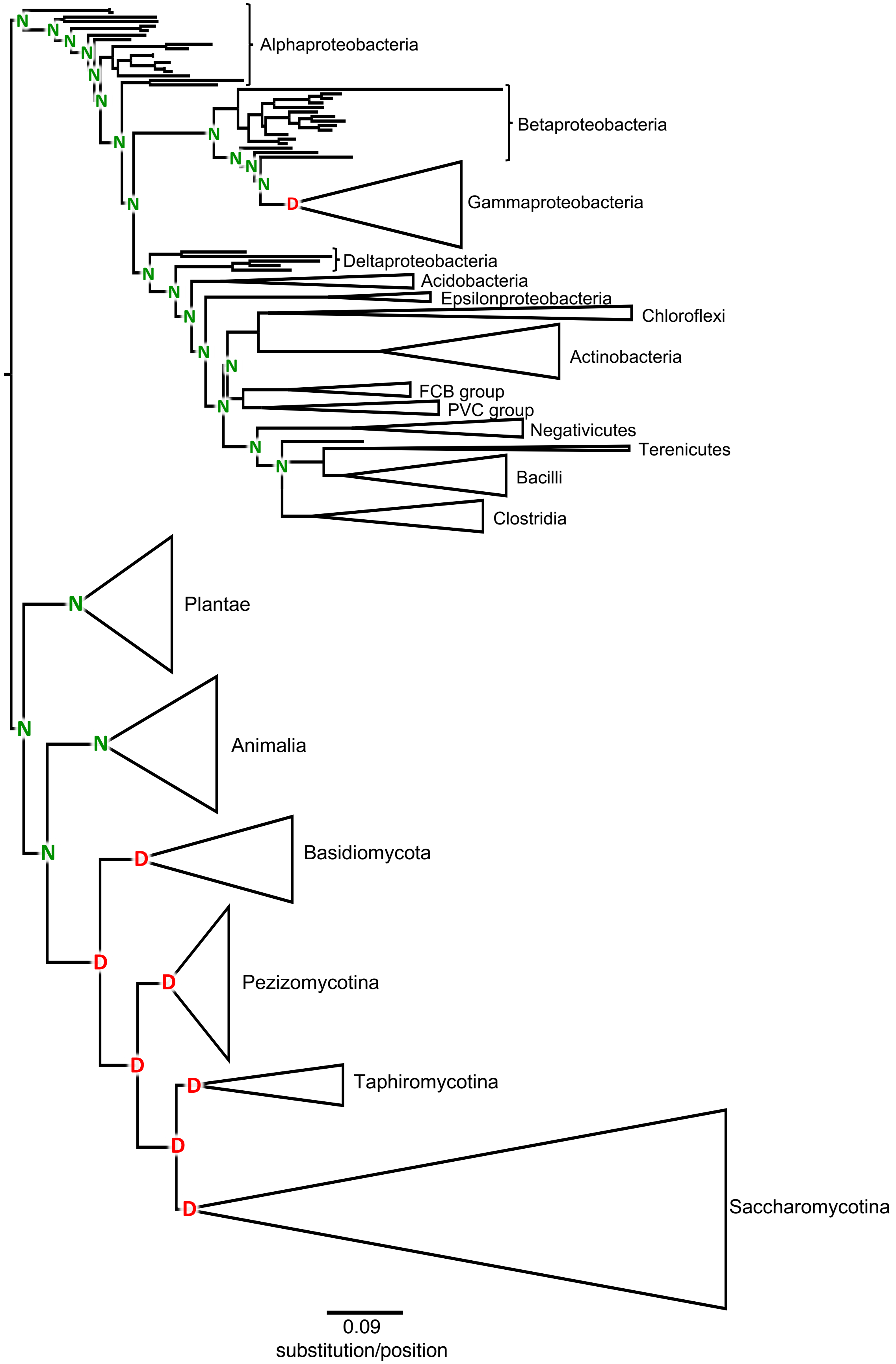


**Supplementary Fig. S6**

Phylogenetic distribution and ancestral reconstruction of aspartate and asparagine residues at the site homologous to $D^{SBD}$ of DnaK and Ssq1; amino acid state is indicated at the nodes of the maximum likelihood phylogeny of DnaK and mitochondrial Hsp70 orthologs from 289 species belonging to the indicated clades. Scale is in amino acid substitutions per position.

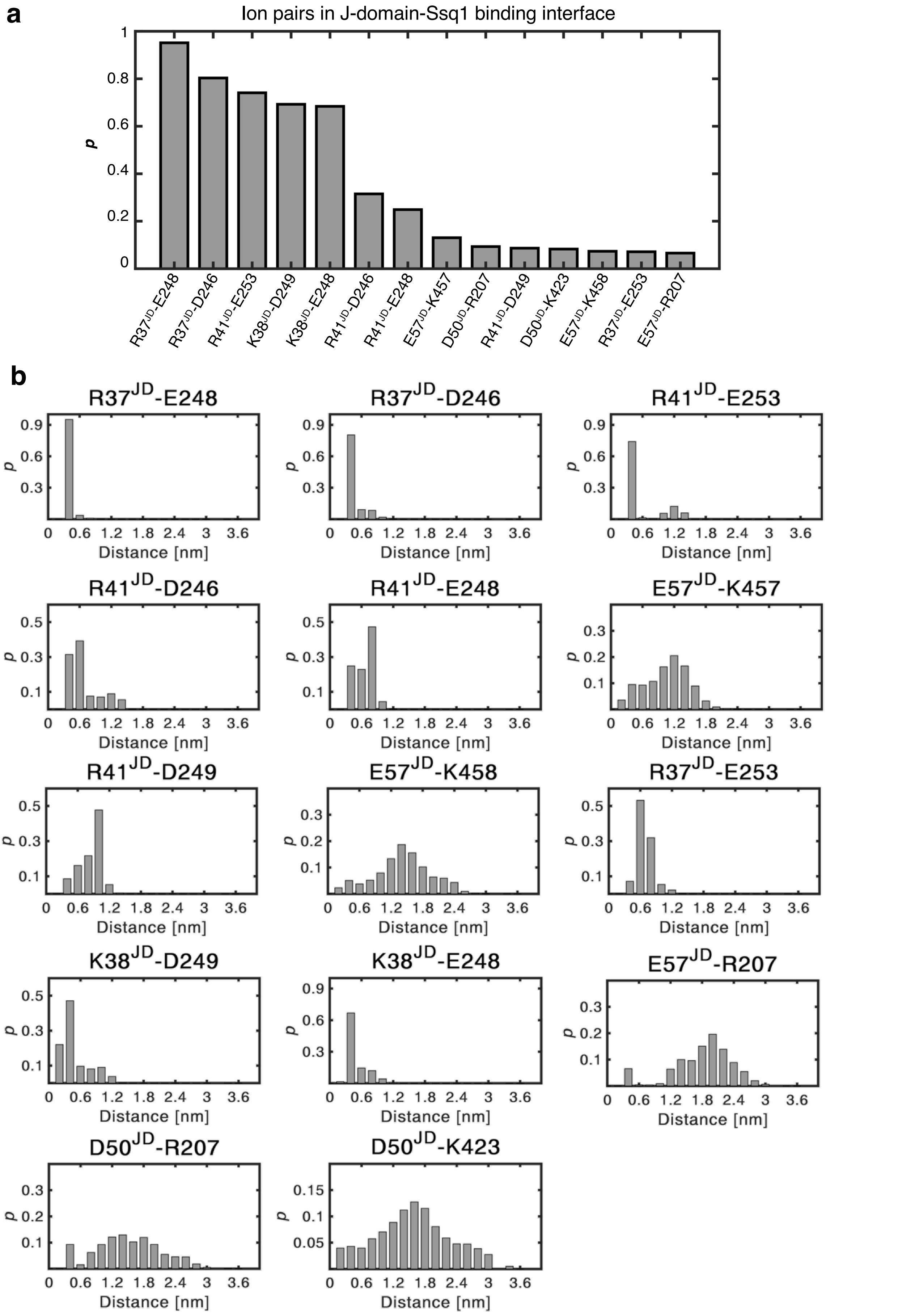


**Supplementary Fig. S7**

(a) Probabilities (p) of the most stable ion pairs across the J-domain/Ssq1 interface, calculated from the 10.5 μs trajectory of the dominant bound state. (b) Distance distributions of the ion pairs shown in (a).

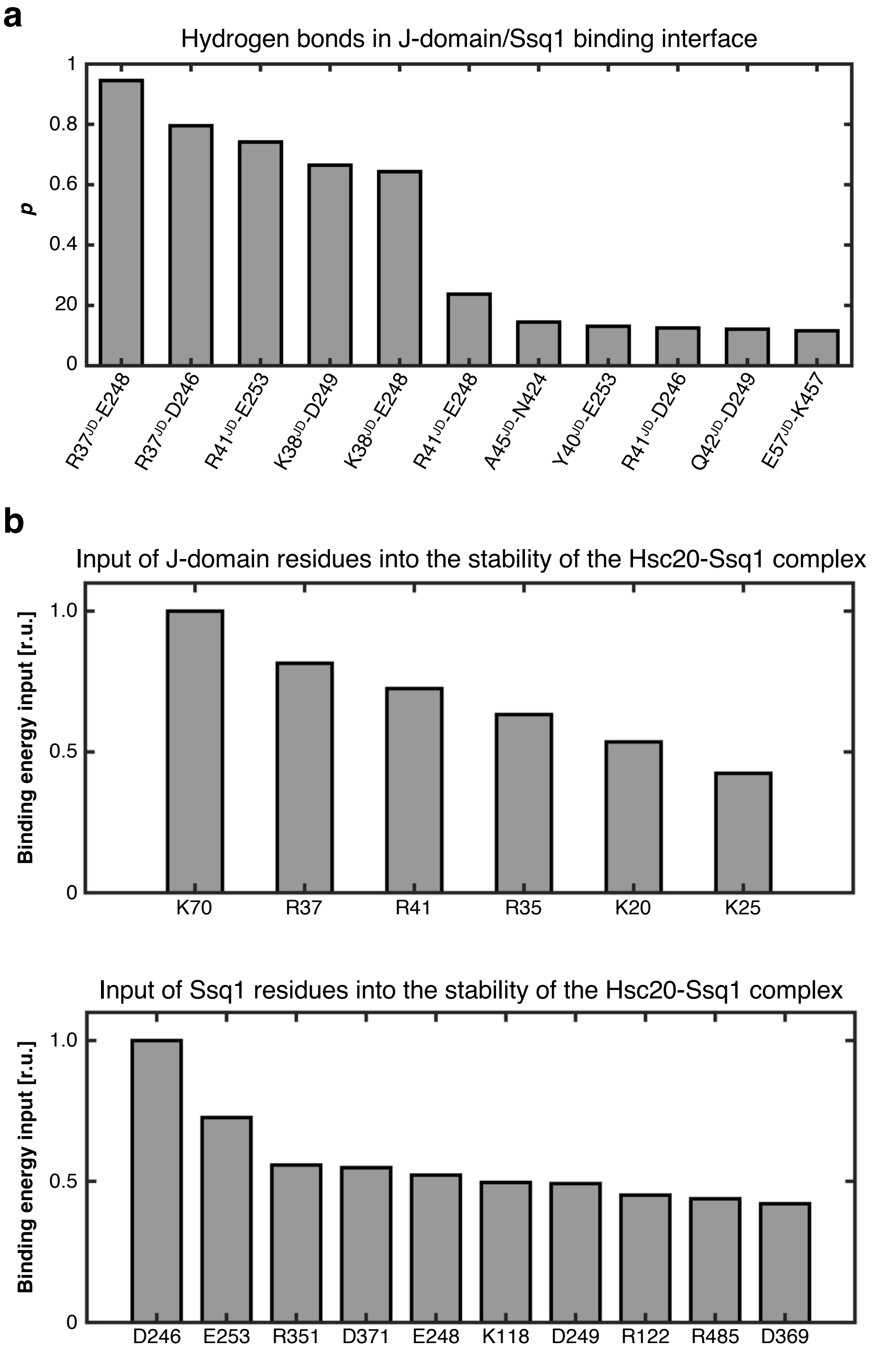


**Supplementary Fig. S8**

(a) Probabilities (p) of the most stable hydrogen bonds across the J-domain/Ssq1 interface, calculated from the 10.5 μs trajectory of the dominant bound state. (b) Relative energy inputs to Hsc20-Ssq1 complex binding energy calculated with MM/PBSA for the residues of the Hsc20 J-domain (top) and Ssq1 (bottom).

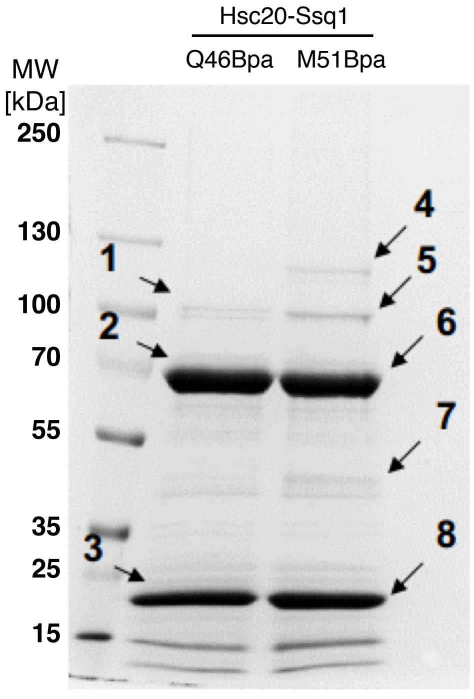


| Band | Protein | MW [Da] | MASCOT Score | emPAI |
|---|---|---|---|---|
| 1 | Ssq1 | 72378 | 8231 | 14.51 |
| | Hsc20 | 21822 | 1658 | 7.14 |
| 2 | Ssq1 | 72378 | 38546 | 40.80 |
| 3 | Hsc20 | 21822 | 10706 | 10.92 |
| | Ssq1 | 72378 | 10259 | 11.28 |
| 4 | Hsc20 | 21822 | 13640 | 16.43 |
| | Ssq1 | 72378 | 2637 | 7.14 |
| 5 | Ssq1 | 72378 | 13350 | 18.48 |
| | Hsc20 | 21822 | 2790 | 7.14 |
| 6 | Ssq1 | 72378 | 34076 | 26.79 |
| 7 | Ssq1 | 72378 | 14297 | 15.44 |
| | Hsc20 | 21822 | 3283 | 7.14 |
| 8 | Hsc20 | 21822 | 9675 | 10.92 |
| | Ssq1 | 72738 | 3798 | 7.16 |

**Supplementary Fig. S9**

Identified products of Hsc20Bpa and Ssq1 WT site-specific UV-crosslinking.

(left) Purified Hsc20 Q46Bpa and M51Bpa variants, were incubated with Ssq1 WT in the presence of ATP, irradiated with UV light and separated by SDS-PAGE. Indicated bands (1-8) were excised and subjected to trypsin digestion followed by LC-MS. Migrations of size standard, in kDa, are indicated. (right) Identification of proteins detected in individual SDS-PAGE bands by Mascot, along with statistical significance of Peptide Mass Fingerprint search (Mascot score) and protein abundance estimation (emPAI).

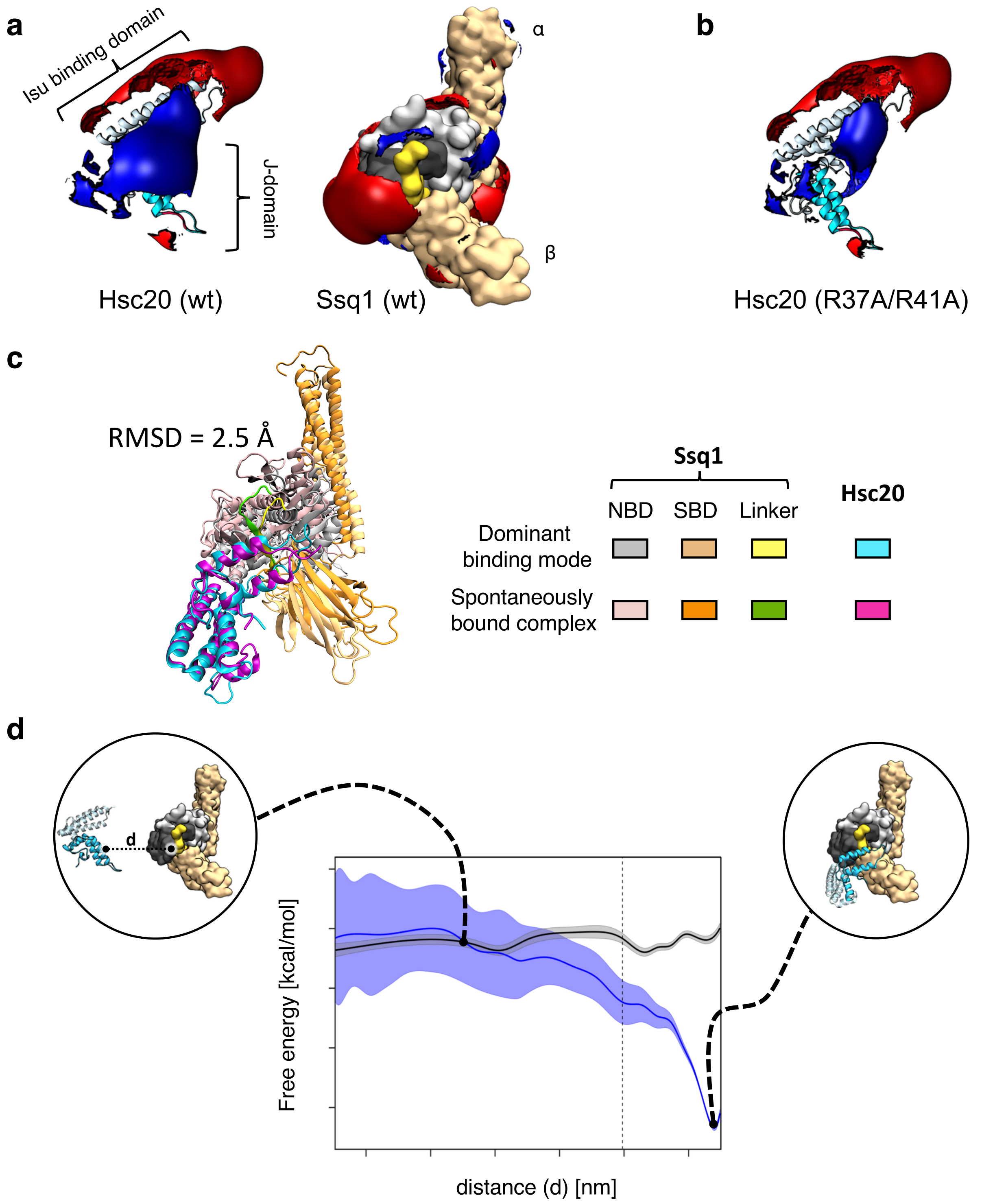


**Supplementary Fig. S10**

Hsc20/Ssq1 recognition is driven by long-range electrostatic forces

(a) Electrostatic isopotential contours at +/- 1kT/e (blue and red, respectively) around Hsc20 and Ssq1 and (b) around Hsc20 double substitution variant R37A,R41A. Coloring of structures as in Fig. 1: Hsc20: J-domain (cyan); Isu binding domain (light blue); Ssq1: SBDα and SBDβ (brown); interdomain linker (yellow); NBD subdomains Ia and IIa in light and dark grey, respectively. (c) Structural alignment of Hsc20-Ssq1 complex obtained by spontaneous binding MD simulations and the dominant bound state obtained by molecular docking/MD simulations. The heavy-atom RMSD for J-domains after aligning NBDs equals to 2.5 Å. (d) Free energy profiles for Hsc20-Ssq1 binding as a function of separation distance, d, between J-domain of Hsc20 and NBD subdomain IIa of Ssq1. Binding of Hsc20 wild-type (WT) (blue line), binding of Hsc20(R37A,R41A) variant (red line); shaded areas show the standard error. The vertical dashed line, demarcating the bund and unbound states, indicates the distance beyond which all specific interactions between the binding partners are lost. Representative structures are shown in circles: unbound state (left), bound state - corresponding to the energy well (right).

**Supplementary Table 1 | Force constants used in Umbrella Sampling for Hsc20-Ssq1 spontaneous binding**

| Hsc20 WT | | Hsc20 R37A R41A | |
|---|---|---|---|
| Distance [nm] | Force constant [kJ/mol * $nm^2$] | Distance [nm] | Force constant [kJ/mol * $nm^2$] |
| 1.2 | 3000 | - | - |
| 1.3 | 3500 | 1.3 | 3500 |
| 1.4 | 4000 | 1.4 | 4000 |
| 1.5 | 3000 | 1.5 | 3000 |
| 1.6 | 3000 | 1.6 | 3000 |
| 1.7 | 2500 | 1.7 | 2500 |
| 1.8 | 2000 | 1.8 | 2000 |
| 2.0 | 1500 | 2.0 | 1500 |
| 2.2 | 1000 | 2.2 | 1000 |
| 2.4 | 1000 | 2.4 | 1000 |
| 2.6 | 1000 | 2.6 | 1000 |
| 2.8 | 1000 | 2.8 | 1000 |
| 3.0 | 1000 | 3.0 | 1000 |
| 3.2 | 1000 | 3.2 | 1000 |
| 3.4 | 500 | 3.4 | 500 |
| 3.6 | 500 | 3.6 | 500 |
| 3.8 | 500 | 3.8 | 500 |
| 4.0 | 500 | 4.0 | 500 |
| 4.2 | 500 | 4.2 | 500 |

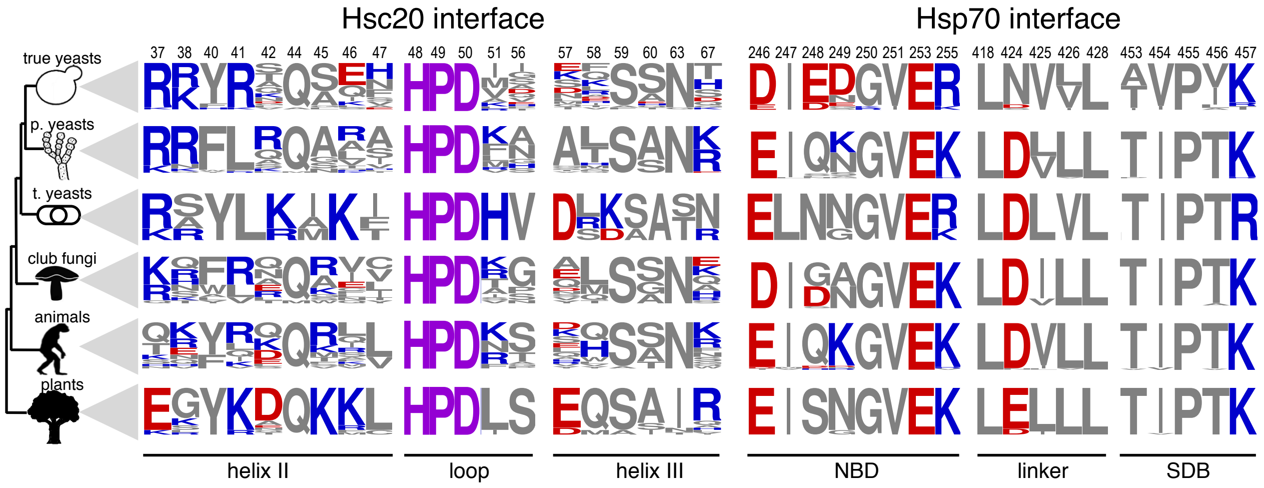


**Supplementary Fig. S11**

Sequence variability of positions that form the Hsc20/Ssq1 binding interfaces (positions shown in green in Fig. 5a) for orthologs of Hsc20 (left) and orthologs of Ssq1 (right) from the following taxonomic groups: Saccharomycotina (true yeasts), Pezizomycotina (p. yeasts), Taphiromycotina (t. yeasts), Basidiomycota (club fungi), Animals (animals) and Plants (plants). Sequence logos represent the amino acid frequency of each interfacial position in the orthologs examined, with positively charged (blue), negatively charged (red) and uncharged (grey) residues. Position numbering is that for *S. cerevisiae*. Phylogenetic relationships among taxonomic groups are depicted on the left.

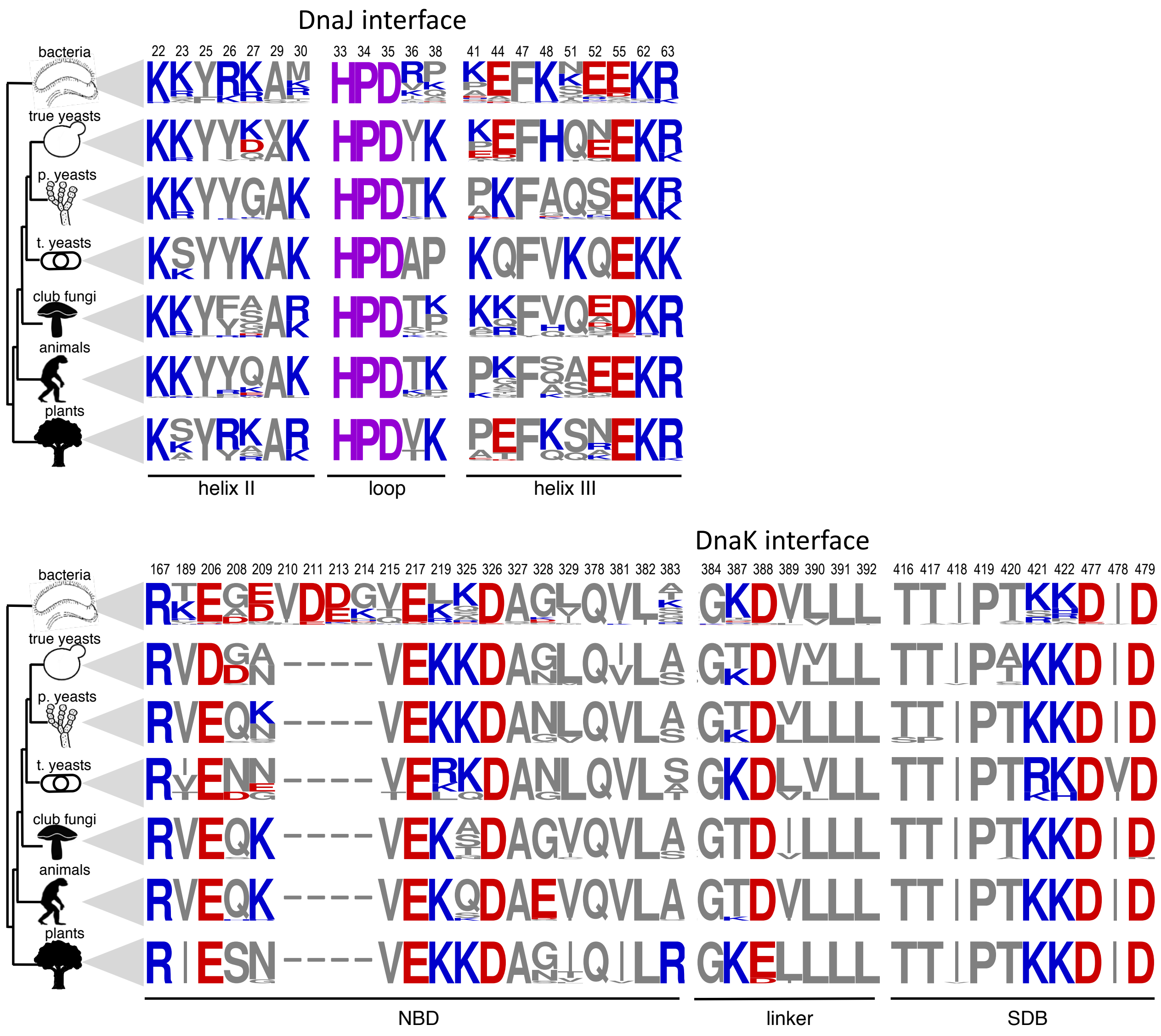


**Supplementary Fig. S12**

Sequence variability of positions that form the DnaJ/DnaK binding interfaces (positions shown in green in Fig. 5a) for orthologs of DnaJ (top) and orthologs of DnaK (bottom) from bacteria and from mitochondria for the following taxonomic groups: Bacteria, Saccharomycotina (true yeasts), Pezizomycotina (p. yeasts), Taphiromycotina (t. yeasts), Basidiomycota (club fungi), Animals (animals) and Plants (plants). Sequence logos represent the amino acid frequency of each interfacial position in the orthologs examined, with positively charged (blue), negatively charged (red) and uncharged (grey) residues. Position numbering is that for *E. coli*. Phylogenetic relationships among taxonomic groups are depicted on the left.

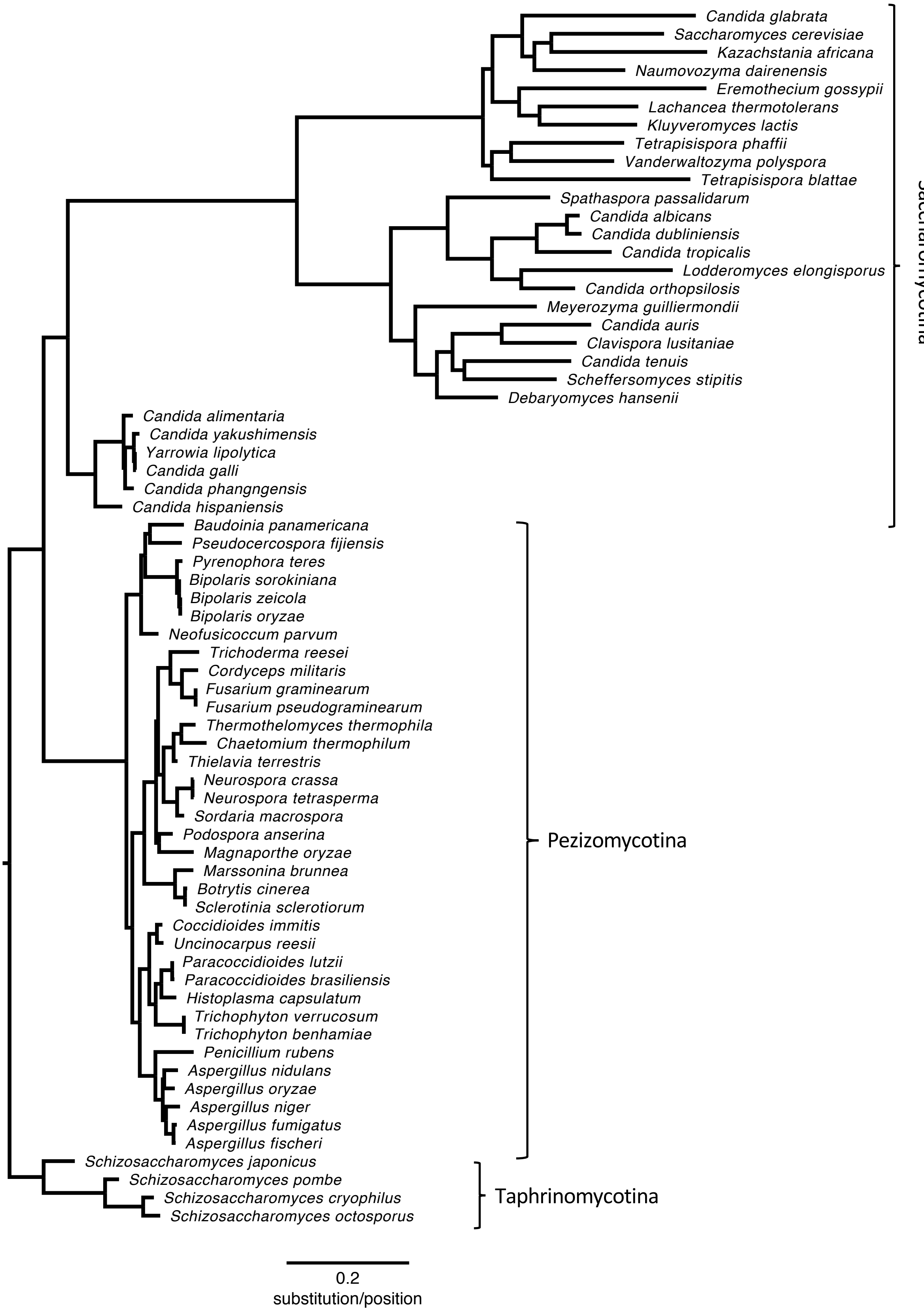


**Supplementary Fig. S13**

Maximum likelihood phylogeny of mitochondrial orthologs of Ssq1 from 67 Ascomycota species used in the sequence variability and co-evolution analyses. Major clades are indicated.

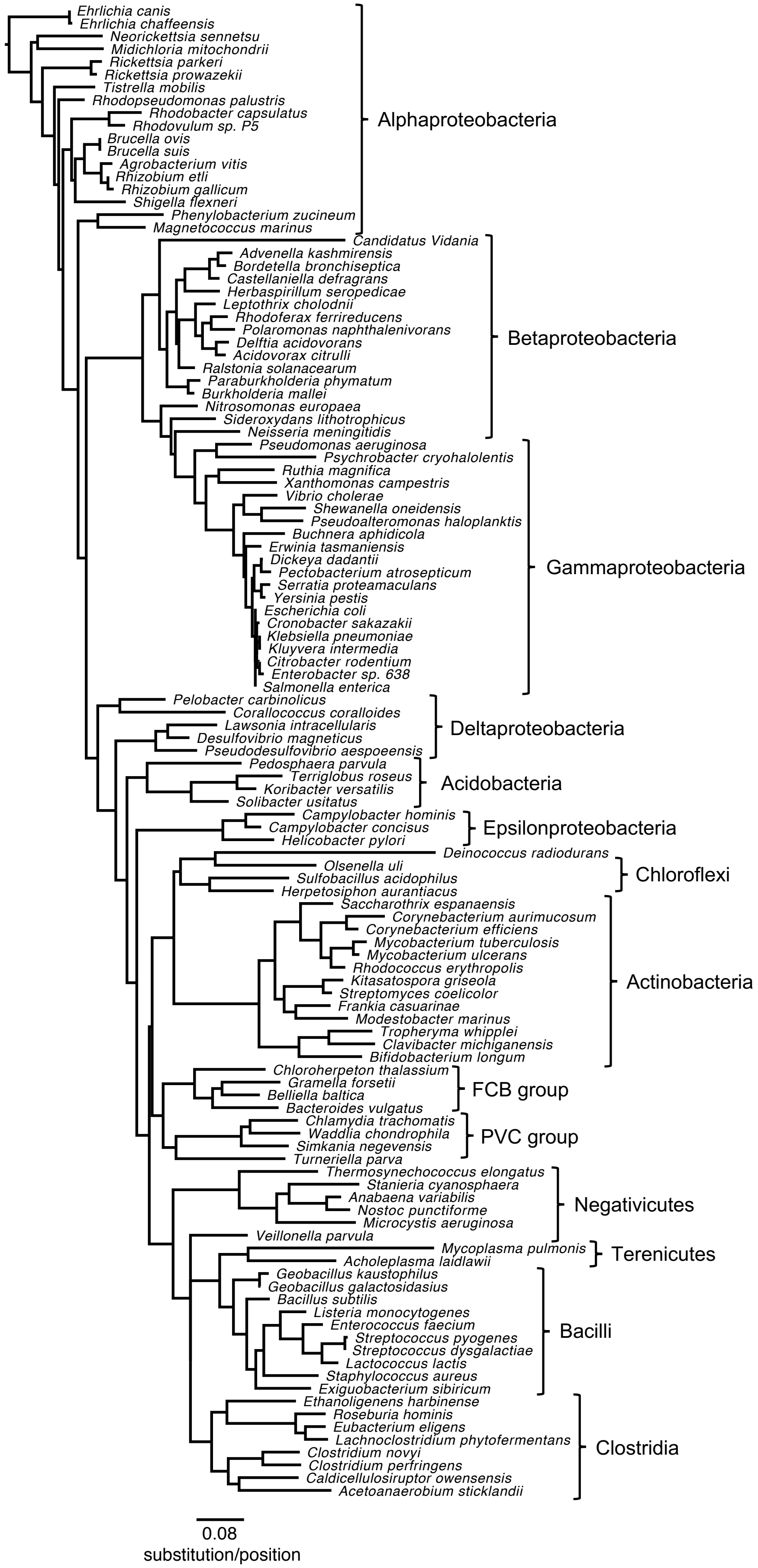


**Supplementary Fig. S14**

Maximum likelihood phylogeny of 117 bacterial DnaK orthologs used in the sequence variability and coevolution analyses. Major clades are indicated.

**Supplementary Table 2 | Statistical support for pairs of coevolving Hsc20/Hsp70 positions from Ascomycota.**

| coevolving positions Hsc20-Hsp70 | lnl Coev | AIC* Coev | lnl M0 | AIC* M0 | ΔAIC | *d/s*** |
|---|---|---|---|---|---|---|
| 38-248 | -118.16 | 240.33 | -134.49 | 270.97 | 30.64 | 8.42 |
| 67-249 | -157.21 | 318.42 | -172.18 | 346.36 | 27.94 | 3.68 |
| 45-249 | -162.84 | 329.69 | -174.65 | 351.30 | 21.61 | 10.00 |
| 42-456 | -171.56 | 347.12 | -182.63 | 367.27 | 20.15 | 5.26 |
| 42-248 | -166.20 | 336.41 | -176.24 | 354.48 | 18.07 | 5.26 |
| 51-249 | -173.68 | 351.36 | -183.12 | 368.23 | 16.88 | 4.00 |
| 38-249 | -165.79 | 335.58 | -175.09 | 352.18 | 16.60 | 5.71 |
| 67-456 | -128.99 | 261.98 | -137.83 | 277.66 | 15.67 | 10.00 |
| 46-248 | -169.43 | 342.87 | -177.65 | 357.31 | 14.44 | 5.26 |
| 42-457 | -164.62 | 333.25 | -172.45 | 346.90 | 13.65 | 10.00 |
| 46-456 | -176.32 | 356.64 | -184.05 | 370.10 | 13.45 | 5.26 |
| 45-248 | -126.84 | 257.69 | -133.90 | 269.81 | 12.12 | 10.00 |
| 42-424 | -164.19 | 332.37 | -170.95 | 343.90 | 11.52 | 5.57 |
| 42-453 | -161.25 | 326.50 | -167.81 | 337.62 | 11.12 | 10.00 |
| 44-248 | -67.08 | 138.17 | -73.13 | 148.26 | 10.09 | 8.42 |
| 67-248 | -125.54 | 255.09 | -131.43 | 264.87 | 9.78 | 17.50 |

* AIC Akaike information criterion
** The *d/s* ratio represents the strength of coevolution between a pair of position in the Coev model. *s* is the rate at which a coevolving pair is replaced by a non-coevolving pair. *d* is the rate at which the pair of positions returns to the coevolving profile. Therefore, *d/s* represents the attraction of pairs of positions to stay within the coevolving profile. *d/s* = 1 indicates lack of coevolution, *d/s* larger than one indicates the strength of coevolution.

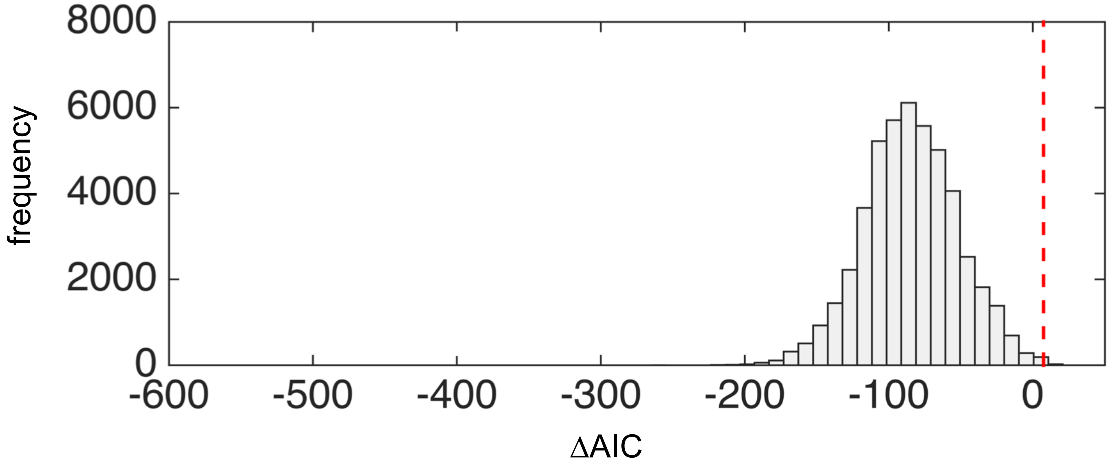


**Supplementary Fig. S15**
ΔAIC ($AIC_{independent}$ - $AIC_{Coev}$) distribution calculated for each pair of positions derived from amino acid sequence of length 33 simulated under the independent evolution model using mitochondrial Hsp70 phylogeny (Fig. S12). The co-evolution acceptance threshold with $p$>0.01 confidence was established as ΔAIC = 7.38 (red dashed line).

**Supplementary Table 3 | Statistical support for pairs of coevolving $DnaJ^{JD}$/DnaK positions from bacteria.**

| coevolving positions $DnaJ^{JD}$-DnaK | lnl Coev | AIC* Coev | lnl M0 | AIC* M0 | ΔAIC | *d/s*** |
|---|---|---|---|---|---|---|
| 51-329 | -281.67 | 567.33 | -297.26 | 596.53 | 29.19 | 10.00 |
| 38-209 | -226.02 | 456.05 | -240.94 | 483.87 | 27.83 | 7.27 |
| 26-209 | -113.05 | 230.09 | -124.43 | 250.86 | 20.77 | 8.42 |
| 23-383 | -327.15 | 658.3 | -337.69 | 677.39 | 19.09 | 10.00 |
| 38-189 | -291.85 | 587.7 | -301.78 | 605.57 | 17.87 | 10.00 |
| 23-189 | -265.11 | 534.22 | -274.94 | 551.87 | 17.65 | 7.27 |
| 36-387 | -225.6 | 455.19 | -235.2 | 472.41 | 17.21 | 10.00 |
| 36-381 | -190.06 | 384.13 | -199.38 | 400.77 | 16.64 | 7.27 |
| 62-189 | -190.51 | 385.02 | -199.7 | 401.4 | 16.38 | 10.00 |
| 38-215 | -215.66 | 435.32 | -224.68 | 451.36 | 16.04 | 10.00 |
| 38-387 | -240.64 | 485.28 | -249.07 | 500.13 | 14.85 | 10.00 |
| 51-189 | -289.54 | 583.08 | -297.96 | 597.93 | 14.85 | 10.00 |
| 52-219 | -205.74 | 415.49 | -213.93 | 429.86 | 14.37 | 10.00 |
| 55-420 | -179.54 | 363.08 | -187.63 | 377.27 | 14.18 | 6.82 |
| 23-329 | -266.16 | 536.32 | -274.24 | 550.47 | 14.15 | 10.00 |
| 51-214 | -251.65 | 507.3 | -259.27 | 520.54 | 13.25 | 7.27 |
| 36-420 | -212.05 | 428.11 | -218.78 | 439.55 | 11.45 | 5.71 |
| 52-387 | -165.03 | 334.05 | -171.5 | 345.00 | 10.95 | 10.00 |
| 52-189 | -218.01 | 440.03 | -224.22 | 450.44 | 10.41 | 10.00 |
| 62-214 | -155.28 | 314.56 | -161.01 | 324.01 | 9.45 | 10.00 |

* AIC Akaike information criterion
** The *d/s* ratio represents the strength of coevolution between a pair of position in the Coev model. *s* is the rate at which a coevolving pair is replaced by a non-coevolving pair. *d* is the rate at which the pair of positions returns to the coevolving profile. Therefore, *d/s* represents the attraction of pairs of positions to stay within the coevolving profile. *d/s* = 1 indicates lack of coevolution, *d/s* larger than one indicates the strength of coevolution.

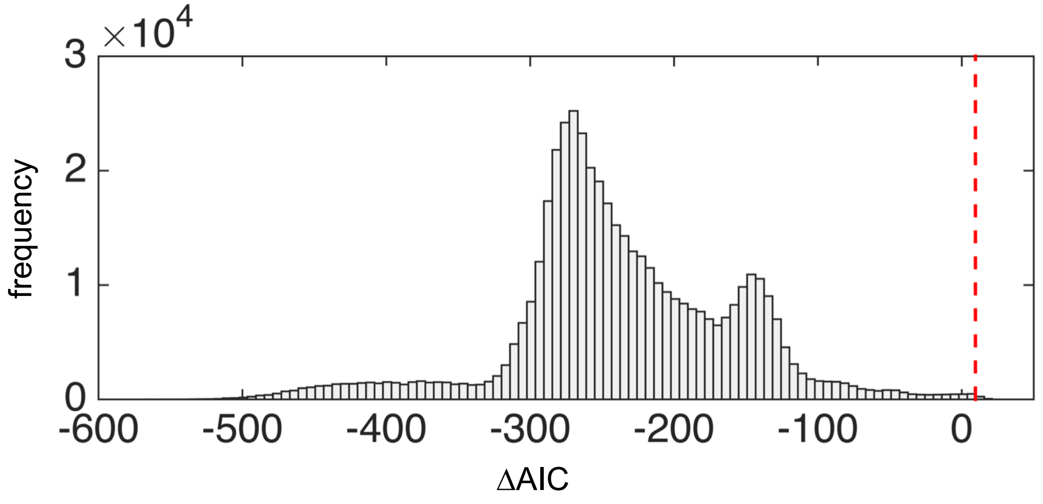


**Supplementary Fig. S16**
ΔAIC ($AIC_{independent}$ - $AIC_{Coev}$) distribution calculated for each pair of positions derived from amino acid sequence of length 54 simulated under the independent evolution model using DnaK phylogeny (Fig. S13). The co-evolution acceptance threshold with $p>0.01$ confidence was established as ΔAIC = 9.473 (red dashed line).
